# Conservation of preparatory neural events regardless of how movement is initiated

**DOI:** 10.1101/189035

**Authors:** Antonio H. Lara, Gamaleldin F. Elsayed, John P. Cunningham, Mark M. Churchland

**Affiliations:** Department of Neuroscience, Columbia University Medical Center, New York, New York, USA. York, USA; Center for Theoretical Neuroscience, Columbia University, New York, New York, USA; David Mahoney Center for Brain and Behavior Research, Columbia University Medical Center, New York, New York, USA; Kavli Institute for Brain Science, Columbia University Medical Center, New York, New York, USA; Grossman Center for the Statistics of Mind, Columbia University Medical Center, New York, New; Department of Statistics, Columbia University, New York, New York, USA

## Abstract

Voluntary movement is believed to be preceded by a preparatory stage. Evidence arises from experiments where a delay separates instruction and execution cues. While this sequence emulates some real-world situations (*e.g.*, swatting a fly upon landing) movements are commonly made at a moment of one’s choosing (reaching for a coffee cup) or are made reactively (intercepting a falling cup). To ascertain whether neural events are conserved across such contexts, we examined motor cortex population-level responses in monkeys when reaches were initiated either after an imposed delay, at a self-chosen time, or reactively with very low latency. We found that the same preparatory and movement-related events were conserved. However, preparation was temporally flexible and could be remarkably brief. Our findings support the existing hypothesis that preparation is an obligatory stage that achieves a consistent state prior to movement. Yet our results reveal that preparation can unfold more rapidly than previously supposed.

## Introduction

Multiple lines of evidence argue that voluntary movement is preceded by a preparatory stage^1-9^. Most fundamentally, the voluntary reaction time (RT, the time between a sensory stimulus and the onset of an evoked movement) is typically longer than expected given afferent and efferent delays, suggesting a time-consuming preparatory process separating sensation from action. RTs typically become shorter when a delay period separates an instruction from a go cue, presumably because preparation has time to complete before the go cue^2,4,8,10^. Neurons in many brain areas - including primary motor cortex (M1) and dorsal premotor cortex (PMd) - respond selectively during the delay^1,4,11-15^. Delay-period activity is predictive of RT variability^4,8,10,16^, and its electrical disruption erases the RT-savings provided by the delay^17^. These observations are consistent with the presence of a neural process that must occur before voluntary movement can be initiated, yet does not itself cause movement and can therefore be completed in advance when circumstances allow.

Although appealing, aspects of the above interpretation remain incomplete, uncertain, or controversial. At the neural level, it remains uncertain whether delay-period activity reflects an essential preparatory stage. It has been argued that it does, and that movement-period neural dynamics are seeded by a preparatory state reflected in delay-period activity^10,18-21^. This hypothesis predicts that there should be a recognizable progression of preparatory and movement-related activity even in the absence of a delay. Determining whether this is true has been challenging. An early study revealed that at least some aspects of delay-period activity are recapitulated without a delay^6^. Yet in a recent study, Ames et al.^22^ found that the neural state during the delay-period was bypassed in the absence of a delay. This result has been variously interpreted as evidence that delay-period activity is beneficial but not essential^22^, that delay-period activity is not preparatory but suppressive^23^, or that the range of acceptable preparatory states is broad^8^. Yet Ames et al. also found that the initial ~50 ms of target-driven response was similar with and without a delay. As in^6^, this finding suggested that early, putatively preparatory aspects of the response may be conserved.

Ambiguity at the neural level is underscored by behavioral results suggesting that preparation may be unnecessary^23^. Preparation has generally been considered to be a time-consuming process that makes a sizeable contribution to the RT^2,4,5,9,10^. Yet under certain circumstance, humans and monkeys display very short RTs^24-26^ despite having no time to prepare in advance. Such RTs appear incompatible with a preparatory stage that consumes considerable time.

We wished to explore the hypothesis that delay-period activity reflects a preparatory process that also occurs when there is no delay, but displays enough temporal flexibility to allow very short RTs. In principle, this hypothesis is readily tested via a straightforward strategy: by exploiting the delay period to identify putatively preparatory neural activity, then inquiring whether and how similar activity is present in other contexts.

Unfortunately, this strategy is difficult or impossible to accomplish at the level of single neurons. Most neurons with delay-period activity also exhibit movement-related activity. Indeed, a neuron that primarily displays delay-period activity for some movements can primarily display movement-related activity for other movements. Thus, in the absence of a delay, any activity that occurs between the go cue and movement onset could be preparatory, movement-related, or some combination of the two. One might hope to make a distinction base on timing: pinpointing a time before which activity is putatively preparatory and after which it is putatively movement-related. However, the nature of such timing is unknown, and is itself one of the subjects of inquiry. To identify putatively preparatory and movement-related patterns, one must therefore move beyond single-neuron responses, and instead focus upon population-level properties.

To this end, we leveraged the recent finding^27^ that delay-period activity and movement-related activity occupy orthogonal subspaces. This allowed us to use responses during a delay to define a putatively-preparatory subspace, and responses during movement to define a putatively-movement subspace. The temporal separation provided by the delay period makes it possible to define these subspaces based on well-separated epochs, avoiding the interval where the transition from preparation to movement is uncertain. We further exploited the empirical fact that these spaces are nearly orthogonal, which allows activity in one subspace to be measured independently of activity in the other subspace. Having identified the relevant subspaces, we then observed activity in those subspaces when movements were initiated without an experimenter-imposed delay. Specifically, monkeys performed reaches in three interleaved contexts. The ‘cue-initiated’ context employed a standard delay period (which was used was to define the spaces), the ‘self-initiated’ context allowed the monkey freedom to decide when to move, and the ‘quasi-automatic’ context required monkeys to reactively intercept a moving target with no time to prepare in advance.

We found that both preparatory-subspace and movement-subspace patterns of neural activity were conserved across contexts. For example, quasi-automatic reaches, although executed with very short RTs, exhibited the same sequence of preparatory and movement-related events as self-initiated reaches. However, the time-course of preparatory-subspace events was remarkably flexible across contexts: preparatory-subspace activity developed slowly in anticipation of self-initiated reaches, but lead movement-subspace activity by only a few tens of milliseconds for quasi-automatic reaches. Our results support the hypothesis that a conserved preparatory process is present regardless of whether there exists an experimenter-imposed delay. However, the temporal flexibility of that process is considerable; it can arise well before movement onset, but can also consume surprisingly little time. In agreement with^24,26^, such brevity argues against the idea that movement preparation necessarily involves time-consuming cognitive or high-level planning processes. It is more likely that preparatory activity plays a straightforward and mechanistic role: initializing the circuits that are about to produce descending movement commands.

## Results

We trained two monkeys (Ba and Ax) to execute reaches in three contexts, which differed regarding how movement initiation is cued, and how much time is allowed for preparation. In all contexts, reaches began at a central touch-point and were made to targets arranged radially in the vertical-horizontal plane. The cue-initiated context (**Fig. 1a**) emulated the standard instructed-delay paradigm: a variable delay period (0-1000 ms) separated target onset from an explicit go cue. We analyzed trials with delays >400 ms; shorter delays were included to encourage timely and consistent preparation. In the self-initiated context (**Fig. 1b**) monkeys were free to reach upon target presentation, but waiting longer yielded proportionally larger rewards up to a limit at 1200 ms. Growing reward size was mirrored by growing target size. Thus, available reward was always cued directly; monkeys did not need to estimate elapsed time. Reward (and target size) ceased growing at the moment of movement onset. In the quasi-automatic context (**Fig. 1c**) the target moved rapidly along a radial path towards the screen’s edge. The target moved as soon as it appeared, and monkeys had to intercept it mid-flight. Trials for the three contexts were randomly interleaved. The color of the central touch-point and target specified the context: red, blue and yellow indicated the cue-initiated, self-initiated and quasi-automatic contexts. Monkeys successfully completed the majority of trials for all three contexts: 93% and 95% of cue-initiated trials (monkey Ba and Ax respectively), 94% and 97% of self-initiated trials, and 93% and 93% of quasi-automatic trials.

**Figure 1.**
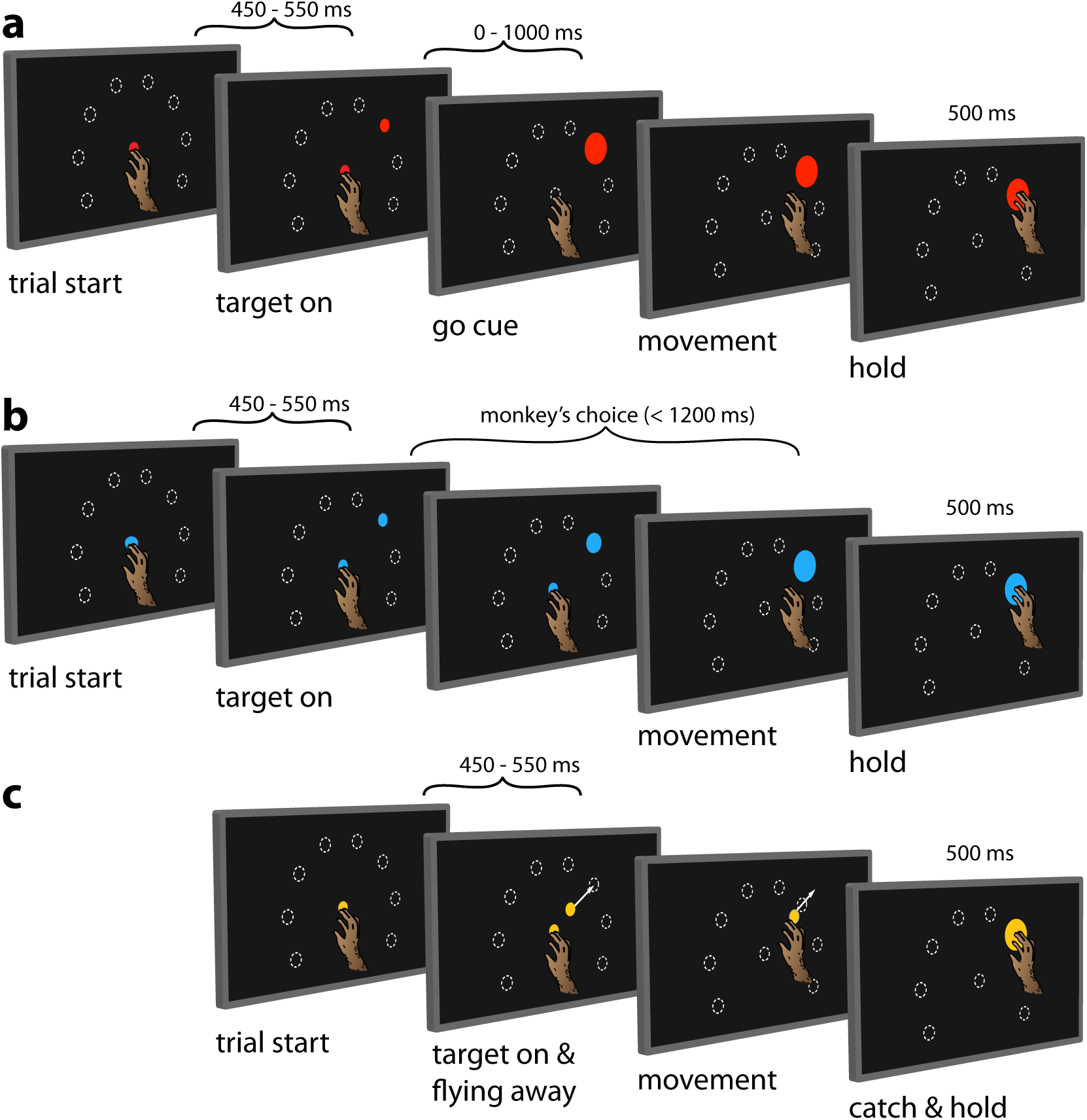
Behavioral task. Monkeys performed the same set of reaches under three initiation contexts**. a)** Cue-initiated context. Trials started when the monkeys touched a red central point on the screen. After a brief delay (450 - 550 ms) a red target appeared in one of eight possible locations (white dashed circles, not visible to the monkey) 130 mm from the touch point. After a variable delay period (0 - 1000ms) the target suddenly increased in size providing the go cue to initiate the reach. **b)** Self-initiated context. Trials began as above, but the central point was blue. Subsequently, a small blue target appeared and gradually grew in size. Monkeys were free to initiate the reach as soon as the target appeared on the screen. However longer waiting times were rewarded with larger amounts of juice. **c)** Quasi-automatic context. The central point was yellow. Yellow targets appeared in one of eight possible locations. The initial appearance of the target was 40 mm from the touch point. Immediately after appearing, the target moved radially outward. Monkeys had to initiate the reach quickly in order to intercept the target before it reached the edge of the screen and disappeared. Targets were intercepted at a location near the location of the targets in the other two tasks (dashed circles).

### Reaction times

RTs were measured based on the moment when hand velocity crossed 1% of its peak (the hand was typically held very steady before movement, allowing this low threshold). In the cue-initiated context, RTs for monkey Ba were 269 ± 50 ms (mean ± s.d.) and RTs for monkey Ax were 251 ± 37 ms (**Fig. 2a,b,** *red traces*). These RTs are on the brisk side of the range reported in prior studies, consistent with the goal of the cue-initiated context: to encourage monkeys to prepare during the delay and reach promptly after the go cue. Despite these brisk RTs, monkeys were almost always successful in waiting for the go cue; reaches during the delay occurred on <1% of trials for both monkeys. Similarly, RTs <150 ms were exceedingly rare, consistent with a go cue whose timing could not be anticipated.

In the self-initiated context, reaches could be made immediately after target onset, but monkeys typically waited at least 600 ms as this garnered a larger reward. Yet monkeys rarely waited until the time of maximum reward (at 1200 ms). We define RT in the self-initiated context as the interval between target and movement onset. Thus, RT has a unified definition across contexts: the time of movement initiation relative to the time when movement was first permitted. Self-initiated RTs were 951 ms ± 132 ms (mean ± s.d.) and 1012 ms ± 106 ms (**Fig. 2a,b**, *blue*). The considerable RT variability in the self-initiated context is unlikely to reflect uncertainty regarding reward size, which was directly conveyed by target size. RT variability presumably reflects the natural tension between a desire for large reward and a desire for immediate reward, with different factors dominating on different trials.

**Figure 2.**
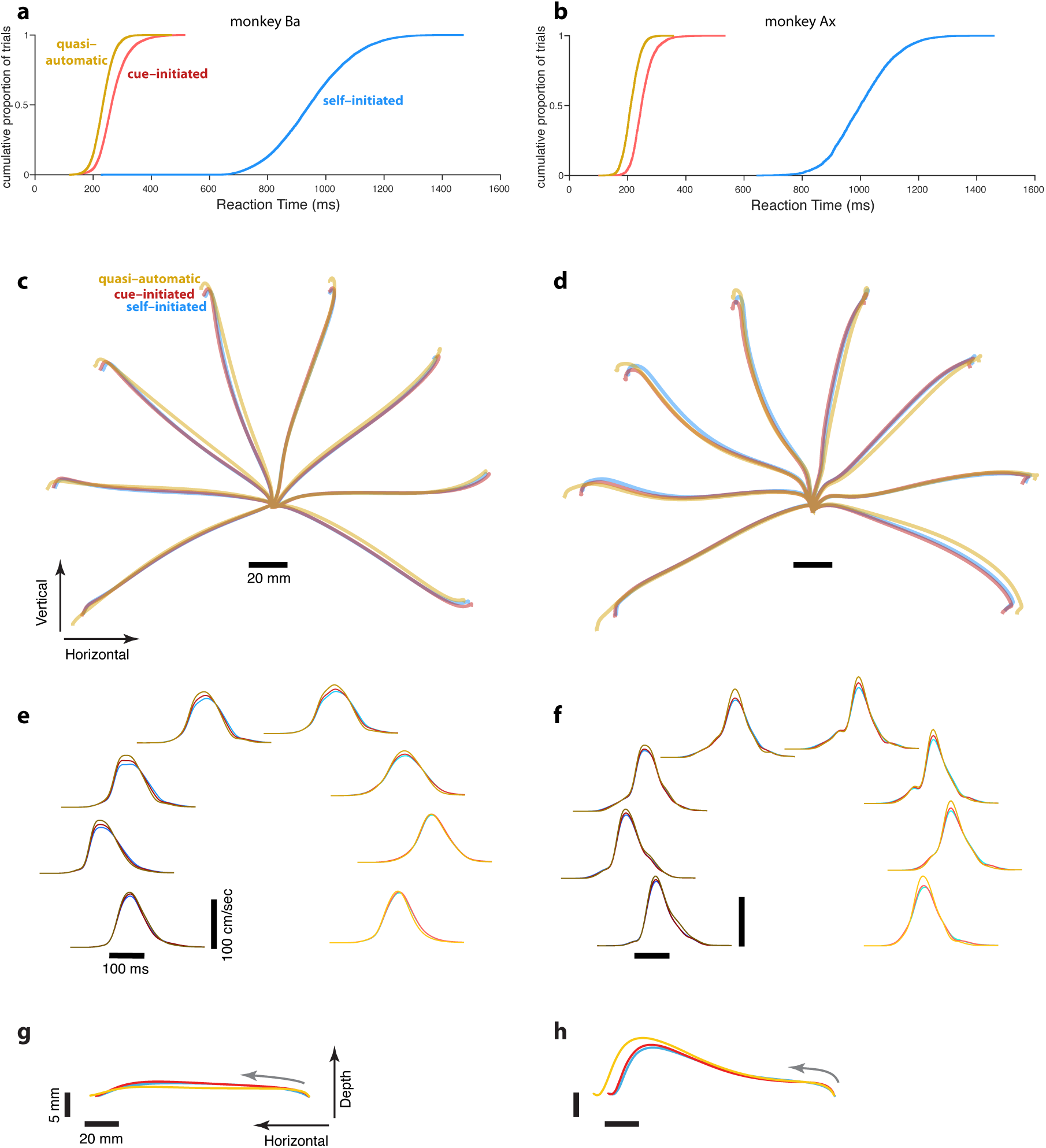
Reaction time distributions and reach kinematics. **a,b)** cumulative reaction time distributions for the three contexts. Trials are pooled across all recordings. **c,d)** Average reach trajectories for the eight targets in the three contexts for monkeys Ba and Ax respectively. **e,f)** Average speed profiles in the three contexts, using the same color coding as above. Additionally, the shade of each line indicates reach direction. This same shade-coding (*light traces* for rightwards reaches and *dark traces* for leftwards reaches) is preserved in subsequent figures. **g,h)** Average reach trajectories for one example reach direction (leftward) with depth shown on an expanded scale to allow closer examination of trajectories in that dimension. Gray arrows indicate direction in which the hand traveled.

The quasi-automatic context evoked particularly short RTs: 221 ± 32 ms and 208 ms ± 27 ms (**Fig. 2a,b,** *yellow*). Notably, RTs were shorter in the quasi-automatic context than in the cue-initiated context (on average by 48 and 43 ms) even though the quasi-automatic context provided no time to prepare in advance. Despite such time pressure, monkeys almost never ‘jumped the gun’. For example, reaches were very rare during the 100 ms interval immediately after target onset (<1% of trials for both monkeys). This was unsurprising: target location was unknown ahead of time, making it impossible to successfully exploit a strategy of anticipatory reaches. Furthermore, any small movement before target onset resulted in an aborted trial, further discouraging anticipatory movements or adjustments. The observed RTs are thus legitimate responses to target onset. Given this, it is notable that RTs could be in the 170-200 ms range. Muscle activity (documented further below) leads this moment by ~80 ms. Thus, muscle activity could begin surprisingly quickly (as little as 90 ms) following target onset.

During training, monkeys showed short RTs immediately upon encountering moving targets in the quasi-automatic context. This observation is consistent with the suggestion that moving targets evoke intercepting movements with an almost innate short latency^26^. For this reason, we refer to such reaches as ‘quasi-automatic’. However, we stress that this term should not be taken to imply that quasi-automatic reaches are necessarily a different class of movement at the level of motor cortex. Whether or not behavior across the three contexts is sub-served by a similar set of preparatory and movement-related neural events is a fundamental question of this study.

### Reach kinematics

To aid comparison of neural activity across contexts, we wished to ensure that any observed differences were not trivially due to differences in the physical reaches themselves. Care was thus taken to ensure that reach trajectories were similar across contexts (**Fig. 2c,d**). For example, for the quasi-automatic context, we adjusted the initial target position and its velocity so that reach extent was similar to that in the other two contexts. The resulting match in kinematics was very good, with only a few slight differences: for many reach directions, quasi-automatic reaches had a slightly greater extent, and a slightly higher corresponding peak velocity (on average 7% and 9% higher relative to cue-initiated movements). It may seem surprising that quasi-automatic reaches did not display dramatically higher velocities compared to the other two contexts. However, we find that monkeys typically reach rapidly even when not required to, presumably out of a desire to obtain reward quickly once the decision to move has been made. Movement duration was indeed similar (~200 ms) across contexts (**Fig. 2e,f**).

We were interested in whether, in the quasi-automatic context, monkeys might begin reaches before they are fully specified^5^. For example, monkeys might initially lift their hand off the screen in a generic fashion and only later adjust their reach towards the target. This could lead to an RT that measures a non-specific response, and would thus not be comparable to RTs measured in the other contexts. To avoid this concern, we measured RT based on the magnitude of the velocity vector in the plane of the task, ignoring depth (distance off the screen). Furthermore, inspection of behavior revealed that monkeys did not adopt the above strategy. Trajectories in depth were very similar across contexts. For example, **Fig. 2g,h** shows example average trajectories in the horizontal / depth plane, with an expanded scale for depth.

Thus, our results confirm that target-directed reaches can be made without a delay period or a long RT^24,26^. We wondered whether some price in accuracy might be paid for such short-latency movements (as might be expected if preparation were hurried). There was indeed a small increase in reach-direction variability in the quasi-automatic context, relative to the cue-initiated context. Assessed via the circular standard deviation^28^ of the initial reach direction, variability was 17% and 11% higher (monkey Ba and Ax; p<0.001 for both). Consistent with the possibility of hurried preparation, quasi-automatic trials with shorter-than-median RTs showed slightly greater variability versus trials with longer-than-median RTs (4.9% and 3.0%; p<0.05 for monkey Ba, NS for monkey Ax). In absolute terms these effects were small – the standard deviation of reach direction was never higher than 4.3 degrees. Still, the quasi-automatic context had slightly more reaches that missed the target: an increase from 1.5% (cue-initiated) to 3.3% (quasi-automatic) for monkey Ba, and from 0.9% to 2.4% for monkey Ax. In summary, very low-latency reaches tended to be slightly less accurate, but were otherwise very similar to reaches in the other two contexts.

### Muscle activity

As with kinematics, muscle activity (EMG) should ideally be similar across contexts to aid comparisons of neural activity. Individual-trial EMG measurements are ‘noisy’ (due to the spiking variability of motoneurons) but the trial-averaged EMG pattern can be accurately estimated by filtering the rectified EMG and averaging across many trials. This process is very similar to that used for the neural data, and employed the same 20 ms standard-deviation Gaussian filter. Trial-averaged EMG patterns were very similar across contexts (**Supplementary Fig. 1**). During movement, the median correlation between the self-initiated and cue-initiated contexts was 0.98 and 0.98 (monkey Ba and Ax respectively; activity compared in a 300 ms window starting 50 ms before movement onset; median taken across muscles). The median correlation between quasi-automatic and cue-initiated contexts was 0.97 and 0.96. We also assessed the strength of each muscle’s selectivity as the standard deviation, across directions, of temporally-averaged EMG. Such selectivity was similar across contexts, but was slightly higher for the quasi-automatic context: by 7.2% and 8.8% (monkey Ba and Ax) relative to the cue-initiated context. This is consistent with the slightly higher extent/velocity for many directions in the quasi-automatic context. Muscle-activity selectivity was very slightly lower in magnitude for the self-initiated context (by 1.9% and 3.0%) relative to the cue-initiated context, consistent with slightly lower velocities for some reach directions (visible upon close inspection of **Fig. 2e,f**).

Many muscles displayed activity while holding the central touch point, before target or movement onset. Such activity was necessary to support the outstretched arm. This affords an opportunity to ask whether putatively preparatory neural activity relates to overt changes in muscle tone. As in many prior studies, we found little change in EMG during intervals when preparation would be expected to occur. We examined EMG during the delay period of the cue-initiated context, and during the interval from target onset until 250 ms before movement onset during the self-initiated context. EMG changed little or not at all following target onset, and showed little or no selectivity for reach direction (**Supplementary Fig. 1**). Specifically, for the cue-initiated context, the across-condition variance of EMG (*i.e.*,selectivity for reach direction) during the delay was 0.76% and 2.8% of the variance during movement. For the self-initiated context, the values were 0.51% and 1.2%. Thus, any preparatory process that might be present in this time-range has minimal direct impact on muscle activity.

### Single-neuron responses during the cue-initiated context

The spikes of well-isolated neurons were recorded from the arm region of motor cortex, including M1 and the immediately adjacent region of PMd (129 and 172 neurons for monkey Ba and Ax). Spike-trains were filtered and trial-averaged to yield an estimate of firing rate as a function of time (**Figure 3** shows examples responses for three neurons). Filtering employed a narrow Gaussian kernel (20 ms SD) to ensure that multi-phasic and temporally structured response features^29^ were not lost. Filtering was performed after concatenating two epochs that were time-locked to target and movement onset, respectively. Filtering the concatenated spike-trains yielded a continuous estimate of rate with no discontinuity. However, it should be kept in mind that the temporal interval between target and movement onset was variable across trials. To ensure the continuous estimate of rate was representative, we chose analysis epochs based on typical behavioral performance. For example, in the quasi-automatic context, epochs were selected such that target onset and movement onset were separated by an interval equal to the mean RT.

**Figure 3.**
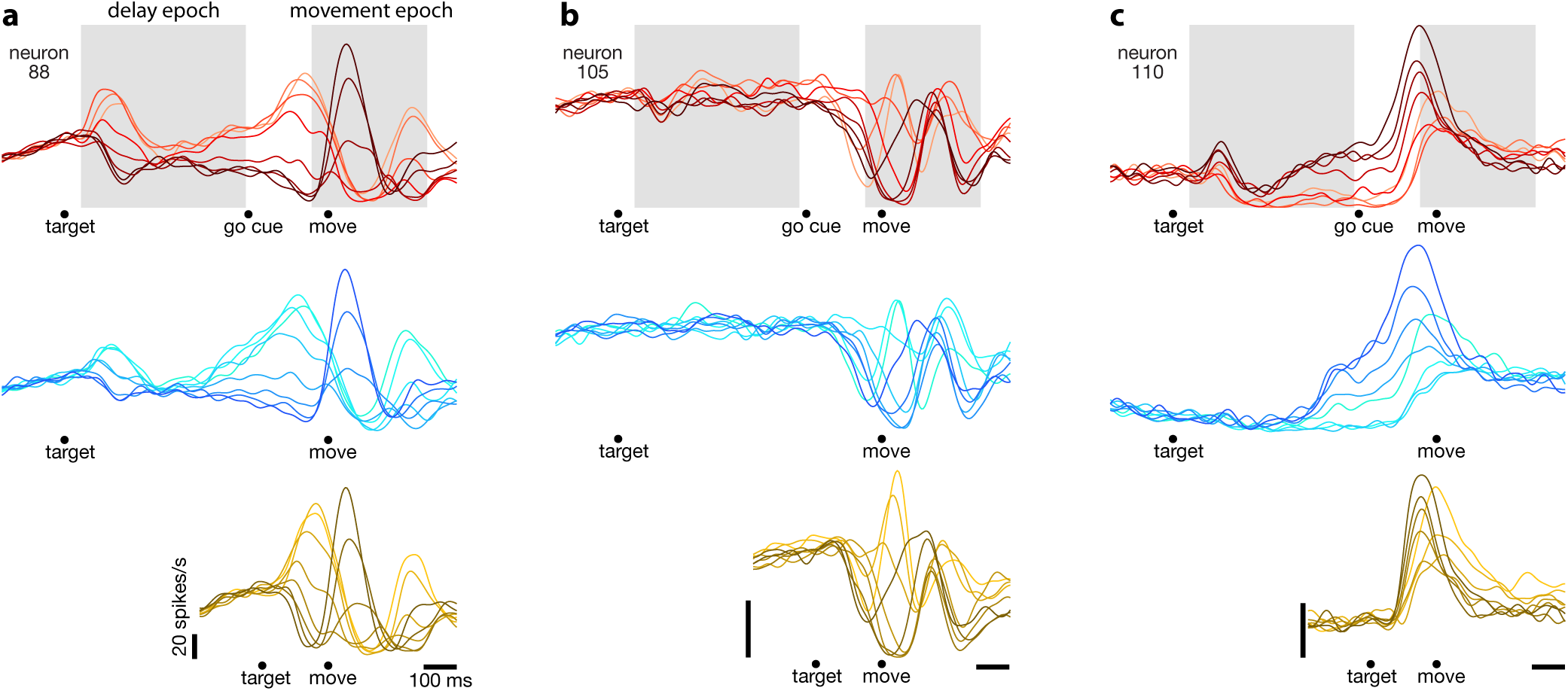
Responses of three example motor cortex neurons. Each column shows the responses of a single neuron for the three initiation contexts. Each trace plots the trial-averaged firing rate for one reach direction (same color scheme as in **Fig. 2**). Gray shaded regions indicate the delay and movement epochs, used to define the preparatory and movement dimensions in subsequent analyses. All traces contain data that was aligned to target onset for the left-hand side of the trace, and to movement onset for the right-hand side of the trace. Individual trials had these two intervals spliced together before filtering and averaging. The sizes of these intervals and the moment of splicing were chosen to imitate the typical timing between target onset and movement onset (keeping in mind that this was variable across trials). For the cue-initiated context, the left-hand side contains data from −200 – 450 ms relative to target onset (only trials with delays >400 ms were analyzed). The right-hand side contains data from −350 ms before movement to 400 ms post movement. The indicated time of the go cue is based on the mean reaction time. For the self-initiated context, spliced averages were computed using the same timing as above, to aid visual comparison. For the quasi-automatic context, the first 150 ms of the response is aligned to the target onset and the subsequent response is aligned to movement onset. Splicing was performed so that the interval from target onset to movement onset matched the mean reaction time. All vertical calibration bars indicate 20 spikes/s.

Many neurons showed activity that varied with reach direction during the delay period of the cue-initiated context (*e.g.*, **Figure 3**, *red traces*). For this context, we defined a 450 ms ‘delay epoch’, beginning 50 ms after target onset. Variation of delay-epoch firing rate with target direction was significant (ANOVA, *p* < 0.05) for the majority of neurons (74/129 and 88/172 for monkey Ba and Ax). We also defined a 300 ms ‘movement epoch’, starting 50 ms before movement onset (just after EMG began to change) and ending just after the hand landed on the target. Variation of movement-epoch firing rate with target direction was also significant for the majority of neurons (116/129 and 144/172).

As in prior studies, delay-period activity suggests a preparatory process. A natural question – relevant to interpretation of activity in the other two contexts – is when putatively preparation-related activity transitions to movement-related activity. Presumably this must happen following the go cue but before movement. Yet the moment of this hypothesized transition is difficult or impossible to determine via inspection of single-neuron responses. For example, the neuron illustrated in **Figure 3a** shows multiple response phases following the go cue, including a peak ~75 ms before movement onset (higher for rightwards reaches; *lighter red traces*) and a subsequent peak during movement (higher for leftwards reaches; *darker red traces*). Should one consider the first peak to be a final strengthening of a preparation-related response, or the beginning of a movement-related response? Is such a distinction even meaningful? These questions are challenging because there is no easily identifiable moment when putatively preparatory activity ends and movement-related activity begins. Similar ambiguity was present even for neurons with simpler response patterns. The neuron illustrated in **Figure 3c** *(red traces)* exhibits activity just before movement that is (approximately) a magnified version of delay-period activity. Again, it is unclear whether such activity reflects the culmination of preparation-related activity, or a movement-related burst.

These uncertainties highlight a known limitation of the instructed-delay paradigm: although activity between target onset and the go cue is suggestive of a preparatory process, it is more challenging to interpret neural events between the go cue and movement onset^6^. One might wish to define events before a specified time as putatively preparatory, and events after that time as movement-related. Yet when that time should be (or even whether the transition happens at a discrete time) is not clear from inspection of single-neuron responses. This creates a problem: if a delay period is necessary to identify activity as putatively preparatory, how can we test whether putatively preparatory events occur in the absence of a delay? This challenge will become particularly relevant when examining responses during the quasi-automatic context.

### Single-neuron responses during the self-initiated versus cue-initiated context

In the self-initiated context, movement initiation is never demanded by an unpredictable go cue. Rather, the monkey chooses when to reach and can potentially anticipate that choice in advance. Given this, how does activity in the self-initiated context relate to that in the cue-initiated context? There exist at least three possibilities. First, if delay-period activity is primarily suppressive, then a similar pattern of activity should be present in the self-initiated context. That pattern should be strongest shortly after target onset (when reaches should be most strongly suppressed because they yield little reward) and should wane as the time of movement approaches. Second, if delay-period activity is primarily preparatory, then during the self-initiated context, a similar pattern should grow with time as movement nears. Any strengthening could be gradual – starting hundreds of milliseconds before movement onset – or rapid – occurring just before movement-related activity. Third, activity in the self-initiated context could look quite unlike activity during the cue-initiated context. Self-initiated movements likely involve a larger role of anterior areas, including the supplementary motor area^30^. If so, the role of motor cortex might be reduced or altered. This could impact pre-movement activity, movement-related activity, or both.

Single-neuron responses followed the second prediction: the patterns of pre-movement activity in the self-initiated context grew with time and came to resemble the patterns of delay-period activity in the cue-initiated context. For example, in **Figure 3a,c**, the ordering of traces ~250 ms before movement onset is similar for the cue-initiated *(red traces)* and self-initiated *(blue traces)* contexts. The pattern of pre-movement activity in the self-initiated context became stronger rather than weaker with time with time. Across all neurons, the median correlation between self-initiated and cue-initiated activity patterns was low during the first 150 ms after target onset (median *r* = 0.39 and *r* = 0.16 for the two monkeys), reflecting the fact that early activity was typically weak in the self-initiated context. This correlation became stronger as movement onset approached (median *r* = 0.86 and *r* = 0.74, using a 150 ms window ending 50 ms before movement onset). During movement, the correlation was quite high (median *r* = 0.92 and *r* = 0.87). Such strengthening of activity agrees with related results in rodent^31^, and is inconsistent with a suppressive process that relaxes to eventually allow movement. These observations will be further quantified by population-level analyses below.

### Single-neuron responses during the quasi-automatic context

Before considering putatively preparatory events, we note that movement-related responses in the quasi-automatic context closely resembled those in the other two contexts. This can be appreciated by visually comparing, across rows in **Figure 3**, activity from just before movement onset until the end of the trial. These example neurons were representative: the median correlation between movement-epoch activity patterns during the quasi-automatic and cue-initiated contexts was 0.85 and 0.85 (monkey Ba and Ax). The primary difference in movement-epoch response patterns was a tendency for some features to be slightly magnified in the quasi-automatic context *(e.g.*, the central peak in **Fig. 3b**). This observation is consistent with the slight increase in reach speed, and with the slight increase in the magnitude of muscle activity. The similarity in movement-related activity across contexts was not a given. Because there may be subcortical contributions to very short-latency movements^25,26^, movement-related cortical activity could have been different or reduced during the quasi-automatic context relative to the other two contexts.

The similarity of movement-related responses makes it sensible to ask whether those responses are preceded by similar patterns of preparatory activity. Is the very first portion of the response in the quasi-automatic context potentially preparatory? Or does that initial response simply constitute the beginning of movement-related activity? We saw no way of addressing this question using individual-neuron analyses. Consider the neurons illustrated **Figure 3a,c**. For both, the very first pattern of activity to emerge in the quasi-automatic context resembled that seen shortly before movement onset in the cue-initiated context. Interpretation thus hinges on whether activity at that time in the cue-initiated context reflects the culmination of preparation or the beginning of movement-related activity. As discussed above, this is difficult or impossible to infer from individual-neuron responses. We therefore turn to analyses that leverage population-level properties.

### Segregating preparatory and movement responses at the population level

We employed a recently developed analytical method to segregate the population response into putatively preparatory and movement-related response patterns. This method leverages the observation that the correlation structure between neurons changes dramatically between delay-period and movement-period epochs^27^. Specifically, the ‘neural dimensions’ that best capture delay-epoch activity do not capture movement-epoch activity, and vice versa. That observation was unexpected; it occurs because neurons with related response properties during preparation become unrelated during movement, something predicted by no existing model^27^. Yet the finding has considerable utility from the standpoint of the present study. Although putatively preparatory and movement-related processes are not separable at the single-neuron level, they are potentially separable at the population level. Our strategy is to use the cue-initiated context to identify a set of neural dimensions that captures putatively preparatory activity and an orthogonal set of dimensions that captures movement-related activity. These dimensions can then be used to examine the population response in the other contexts. If delay-period activity indeed reflects preparation, then dimensions that capture delay-period activity in the cue-initiated context may similarly capture preparatory processing in the other two contexts. Alternatively, if delay-period activity reflects some non-preparatory process (*e.g.*, suppression) or a process specific to the presence of an experimenter-imposed delay, then the dimensions that captured delay-period activity will either not capture activity during the other two contexts, or will capture activity with structure that is very different from that observed during the delay.

Based on neural responses during the cue-initiated context only, we isolated twelve preparatory dimensions, collectively the ‘preparatory subspace’. The preparatory subspace captured 80% of firing rate variance *(i.e.*, firing-rate structure across all neurons) during the delay epoch, but only 3% of variance during the movement epoch. We isolated twelve movement dimensions (collectively the ‘movement subspace’) which together captured 85% of variance during the movement epoch, but only 3% of variance during the delay epoch. The above percentages are for monkey Ba and were similar for monkey Ax (72% versus 4% and 83% versus 3%). The ability to achieve this near-perfect segregation is not a general feature of neural data. It is a consequence of the dramatic change in covariance between the delay and movement epochs.

We projected the population response, for the cue-initiated context, onto the preparatory and movement dimensions. This revealed putatively preparatory and movement-related activity patterns (**Figure 4**, *middle*). Each projection is a weighted sum of single-neuron responses (weights are the elements of the vector defining the dimension). Yet unlike a generic linear readout, weights were optimized to capture response structure. Projections are thus not only readouts but also building blocks of single-neuron responses, much as for principal component analysis. Indeed, the dimensions we found span a space similar to the top principal components. Using these building blocks, it becomes possible to estimate putatively preparatory and movement-related contributions to each neuron’s response. For example, the response of neuron 88 (**Fig. 4**, *rightmost column)* is accurately approximated as the sum of a preparatory-subspace pattern (a weighted sum of preparatory projections, *orange)* and a movement-subspace pattern (a weighted sum of movement projections, *purple*). The reconstruction is not perfect – the pattern in **Figure 4** *(right)* differs slightly from the true response in **Figure 3a** – but is quite good (*R*^2^ = 0.93). High reconstruction accuracy reflects the high proportion of firing-rate variance captured. Do the dimensions found using the cue-initiated context – in particular the preparatory dimensions – similarly capture variance during the other two contexts?

**Figure 4.**
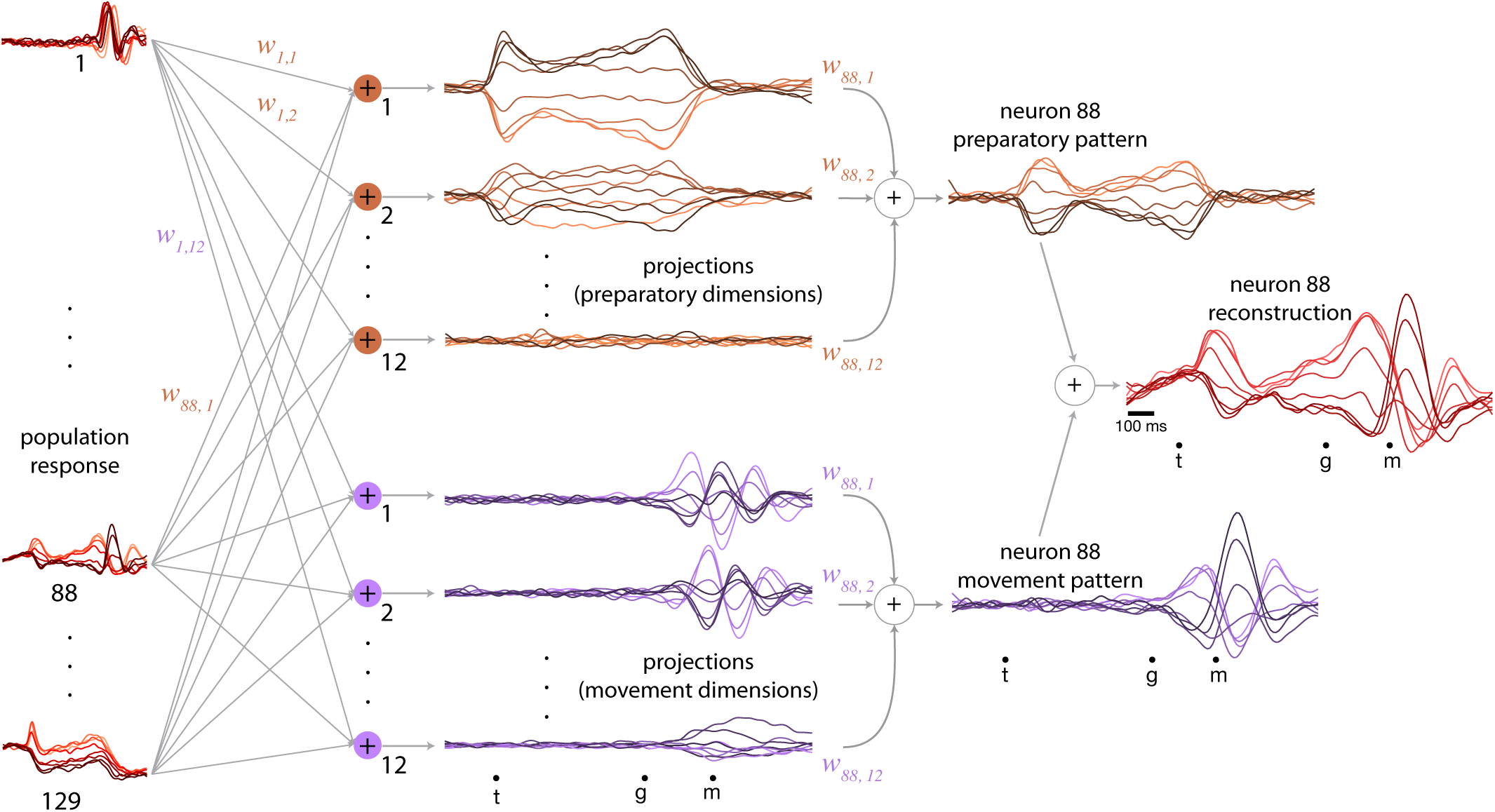
Preparatory and movement contributions to neuronal responses. The population of neural responses *(left column)* can be linearly ‘decode d’ into preparatory and movement-related projections *(middle column). **w_nk_*** is the decode weight from the ***n^th^*** neuron to the /c^th^projectio, and the collection of **w._fe_** such weights i s the *k^th^neural* dimension. Empirically, it is possible to find orthogonal dimensions that segregate preparatory and movement-related response patterns: one set of projections shows structure only during preparation *(orange traces in middle column)*, while the other shows structure only around the time of the movement *(purple traces in middle column*). Because dimensions are chosen to maximally capture data variance (*i.e.*, the structure of firing rates), individual-neuron responses can be reconstructed from the projections. Reconstruction employs the same weights that defined the projections (*e.g.*, if neuron 88 contributed to the first preparatory projection with weight **w_88,1_**, then that first preparatory projection contributes to the reconstruction of neuron 88 with the same weight). The weighted sum of the preparatory projections yields that neuron’s ‘preparatory pattern’ *(orange traces in right column)* and the weighted sum of movement projections yields that neuron’s ‘movement pattern’ *(purple traces in right column*). The full reconstruction is the sum of these two patterns (which describe the tuned aspects of the neuron’s response) plus a time-varying mean that captures any untuned trends in the overall mean firing rate with time (not shown). The success of the approximation can be appreciated by comparison with the true response in Fig. 3a.

### Reconstruction of neural responses across contexts

The dimensions found using the cue-initiated context continued to capture a high percentage of response structure in the self-initiated and quasi-automatic contexts. In the self-initiated context, the total variance captured by both preparatory and movement subspaces was 89% and 84% (monkey Ba and Ax) of that in the cue-initiated context. In the quasi-automatic context, the variance captured was 87% and 84% of that in the cue-initiated context. The high percentage of captured variance is reflected in accurate reconstruction of neural responses in the self-initiated and quasi-automated contexts. For example, the response of neuron 88 (**Fig. 3a**) was accurately reconstructed not only in the cue-initiated context (**Fig. 4**, *right*) but also in the self-initiated (**Fig. 5a**) and quasi-automatic (**Fig. 5b**) contexts.

**Figure 5.**
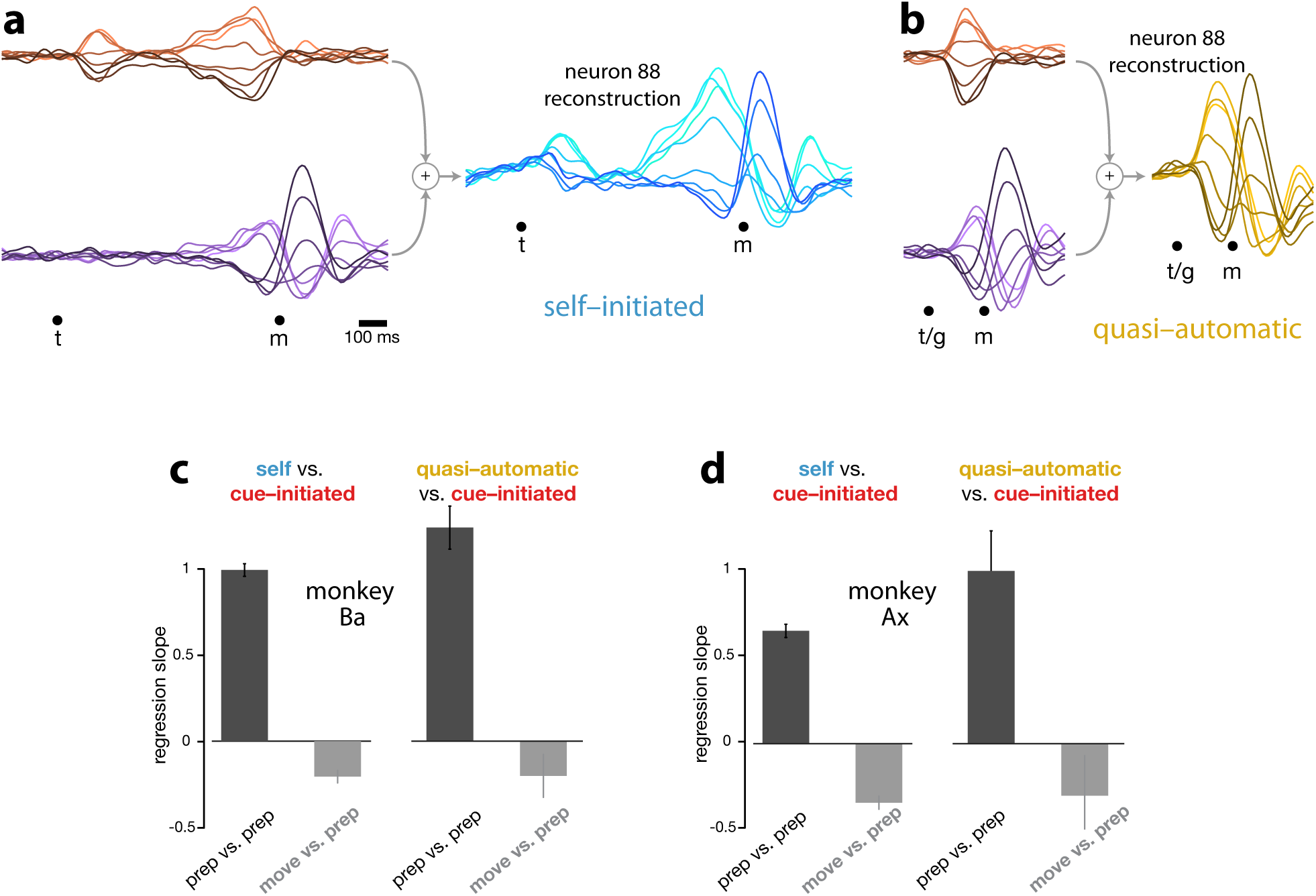
Reconstruction of single-neuron responses in the self-initiated and quasi-automatic contexts, using preparatory and movement dimensions found for the cue-initiated context. **a)**. Example reconstruction for the self-initiated context. The population response was projected onto the preparatory and movement dimensions (found using the cue-initiated context). We then used those projections to reconstruct the preparatory *(orange)* and movement *(purple)* patterns that contributed to each neuron’s response. These patterns are shown for neuron 88. The full reconstruction *(blue)* is the sum of the preparatory and movement-related patterns, plus the across-condition mean response (which has no tuning, but captures the overall mean rate at each time). This reconstructed response can be compared with the true response in Fig. 3a. **b)**. As in **a**, but for the quasi-automatic context. The reconstructed response *(yellow)* can be compared with the true response in Fig. 3a. **c)**. Summary of the degree to which a given neuron’s preparatory-subspace patterns were similar across contexts. Data are for monkey Ba. For each neuron, we took the value of the preparatory subspace pattern for each direction 100 ms before movement onset. These values form a vector. To compute the similarity of the vector for the self-initiated context with that for the cue-initiated context, we regressed the former versus the latter and took the slope. The same was done for the quasi-automatic context versus the cue-initiated context. Dark bars show the average slope across neurons +/- SEM. As a comparison, we repeated this analysis but regressed the movement pattern during the self-initiated and quasi-automatic contexts versus the preparatory pattern during the cue-initiated context *(light gray bars*). The movement pattern was assessed 150 ms after movement onset. **d)**. As in **c**, but for monkey Ax.

These successful reconstructions involved contributions from both preparatory and movement subspaces. For example, the reconstruction for neuron 88 included a robust preparatory-subspace pattern during the self-initiated context (**Fig. 5a**, *orange)* and a short-lived but strong preparatory-subspace pattern during the quasi-automatic context (**Fig. 5b**, *orange*). Within this preparatory-subspace patterns, the ordering of conditions was similar across contexts: for neuron 88 the pattern was most positive for rightwards reaches *(light traces)* and most negative for leftwards reaches *(dark traces*). To facilitate quantitative comparison, for each neuron we measured the preparatory pattern 100 ms before movement onset, yielding a vector with one value per direction. This vector captures the directionality of the preparatory pattern. To assess whether directionality was similar across contexts, for each neuron we regressed the preparatory patterns for the self-initiated and quasi-automatic contexts against that observed for the cue-initiated context. If the two patterns are the same, then regressing one versus the other will yield a slope of one. In contrast, an average slope of zero would indicate no consistent relationship between the preparatory patterns across contexts.

When comparing self-initiated and cue-initiated contexts, slopes were strongly positive (**Fig. 5c, d** *black bars in left subpanel*). When comparing quasi-automatic and cue-initiated contexts, slopes were again strongly positive (**Fig. 5c,d** *dark bars in right subpanel*). For monkey Ba, the slope was slightly greater than unity, consistent with preparatory-subspace activity being slightly stronger in the quasi-automatic context. A potential concern is that this strong similarity might not be specific to the preparatory pattern. For example, if neurons have similar directionality at all times, then similarity would be high even when comparing a preparatory pattern in one context and a movement pattern in another context. This was not the case: there was no consistent relationship between the preparatory-subspace contribution in the cue-initiated context and the movement-subspace contribution (assessed 150 ms after movement onset) in the other two contexts (**Fig. 5c,d**, *gray bars*). In summary, the dimensions that captured delay-period activity also made strong contributions to firing rates during all three contexts, and had a similar pattern across all three contexts. The nature of that pattern will be further investigated below.

### Temporal evolution of responses in preparatory and movement dimensions

If preparatory-subspace activity is truly preparatory, then it should exhibit a time-course consistent with that role. Is this true across all three contexts? For each time, we measured the across-condition variance (the strength of selectivity) of preparatory-subspace activity. This variance reflects the size of the envelope describing the *orange* patterns in **Figure 4**. We refer to this measure as the ‘preparatory-subspace occupancy’. An important question is whether putatively preparatory events consistently precede movement-related events. We therefore similarly computed the movement-subspace occupancy. Movement-subspace occupancy (**Fig. 6**, *purple traces)* had a similar time-course across contexts: it was negligible until ~110 ms before movement onset and reached a peak just after movement onset (the peak occurred between 30 and 80 ms after movement onset across both monkeys and all contexts).

**Figure 6.**
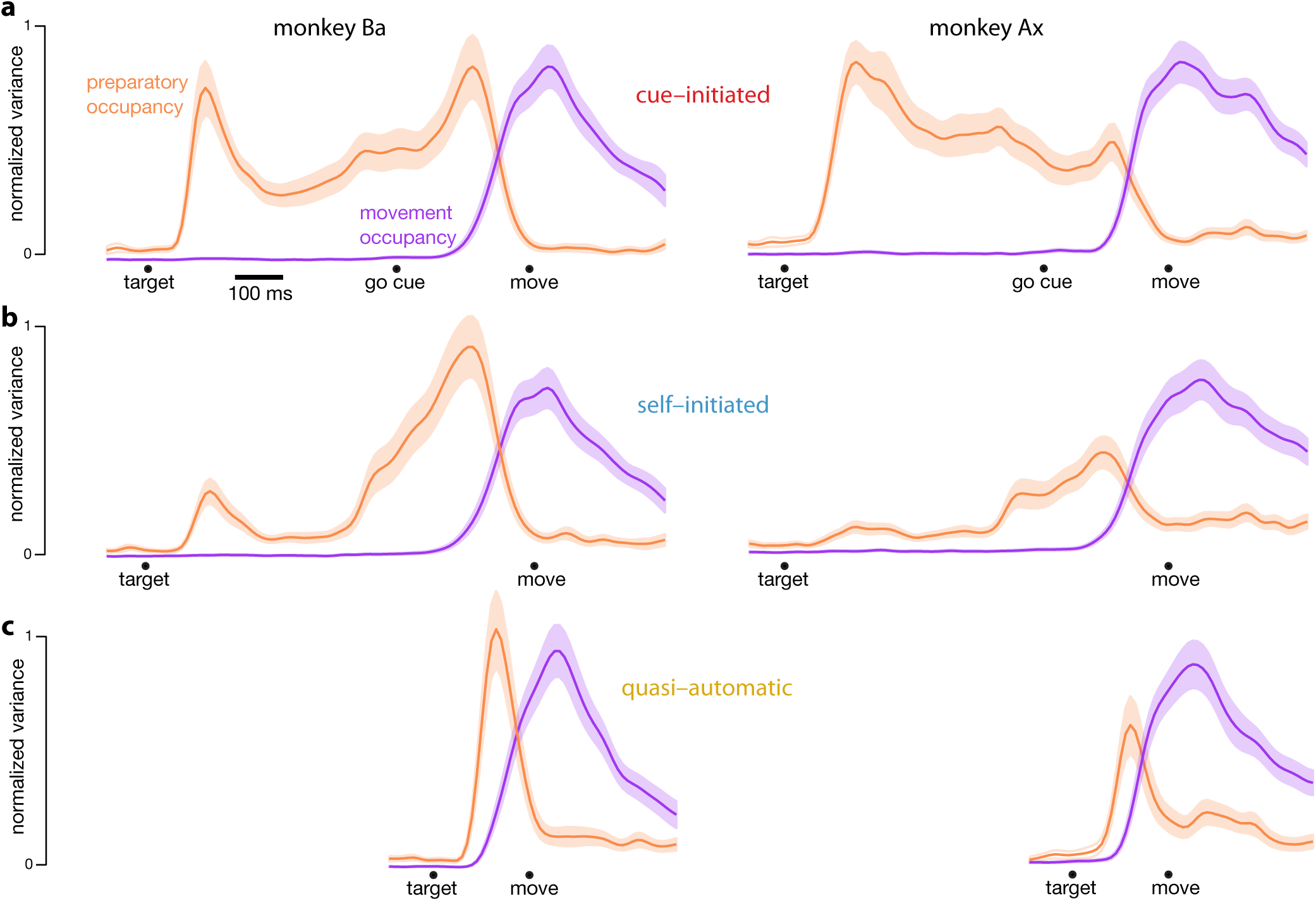
Preparatory and movement-subspace occupancy. **a)** Preparatory and movement subspace occupancy for the cue-initiated context. The two columns show results for monkeys Ba and Ax. Occupancy was calculated as the sum of the across-condition variance in the preparatory and movement dimensions respectively. Preparatory-subspace occupancy, across all three contexts, was normalized by the highest value attained in the cue-initiated context. Movement-subspace occupancy was similarly normalized. The shaded region denotes the standard deviation of the sampling error (equivalent to the standard error) computed via bootstrap *(methods*). **b,c**) Occupancy during the self-initiated and quasi-automatic contexts respectively.

Preparatory-subspace occupancy (**Fig. 6**, *orange traces*) followed a very different time course for each context. In the cue-initiated context, there was an initial rapid rise that was sustained (at a lower level) throughout the delay period. Preparatory-subspace activity then declined rapidly just before movement onset, reaching baseline levels around the time the reach began. It is worth stressing that, by construction, preparatory subspace occupancy is high during the delay period of the cue-initiated context. However, no further structure is imposed; occupancy could have declined following the go cue, could have stayed the same, or could have become stronger.

In the self-initiated context, the rise in preparatory subspace occupancy following target onset was weaker (monkey Ba) or much weaker (monkey Ax) than for the cue-initiated context. Preparatory-subspace occupancy remained weak from 200-400 ms after target onset. Occupancy then grew as movement approached, and reached a peak before movement onset (at 120 ms and 160 ms for monkey Ba and Ax respectively). Occupancy at that time was then similar to occupancy in the cue-initiated context at the same time (slightly greater for monkey Ba and slightly smaller for monkey Ax). These observations concur with the hypothesis that, in the self-initiated context, monkeys do not consistently prepare their reach immediately following target onset, but instead wait until nearer the time they choose to move. Whether the ramp of increasing occupancy reflects ramping on individual trials cannot be inferred from the present data. It is equally plausible that preparation has a sudden onset that is variable relative to movement onset, resulting in a ramp in the averaged data.

In the quasi-automatic context, preparatory-subspace occupancy rose rapidly following target onset and was short-lived: occupancy peaked 70 ms and 80 ms (monkey Ba and Ax respectively) before movement onset and then declined. The magnitude of this peak in preparatory-subspace occupancy was similar to, but slightly higher than, the peak observed in the other two contexts just before movement onset. These observations are consistent with the hypothesis that a preparatory stage is present even for low-latency intercepting reaches. However, preparation appears to be very rapid: preparatory subspace occupancy precedes movement-subspace occupancy by only a few tens of milliseconds: 33 ms for monkey Ba and 42 ms for monkey Ax. (Latency was measured as the time occupancy crossed a 10% threshold, and a shorter filter was used to minimize the influence of filtering on latency, see *methods*).

Comparing between monkeys, there was one obvious difference in the time-course of preparatory subspace occupancy. For monkey Ax, the initial, post-target peak during the cue-initiated context was larger than at any other time, for any context. (Occupancy is plotted in normalized form, and thus the large initial peak for the cue-initiated context necessarily means that all other peaks are plotted with values below unity.) In contrast, for monkey Ba, the post-target and pre-movement peaks in the cue-initiated context were closer in magnitude. When comparing the peak just before movement across contexts, the two monkeys were more similar. For both monkeys, the pre-movement peak in preparatory subspace occupancy was a similar size across contexts, and was slightly larger for the quasi-initiated context. We now ask whether the events within that subspace are conserved across contexts.

### Preparatory and movement events in state space

**Figure 7** plots ‘snapshots’ of projections onto a two-dimensional preparatory subspace (***a***) and a two-dimensional movement subspace (***b***). (**Supplementary Fig. 2** shows similar snapshots for monkey Ax). Within each snapshot, each trace plots the evolution of the neural state for one reach direction over a 150 ms period, beginning at the indicated time. For this task, preparatory-subspace activity was quite low-dimensional: the first two dimensions captured much more variance than subsequent dimensions. *E.g.*, for monkey Ba, the third preparatory dimension captured only 14% as much variance as the first. Thus, the preparatory subspace projections in **Figure 7a** give a reasonably complete view of the preparatory state, and subsequent quantification is based on those dimensions. Movement subspace activity was considerably higher dimensional: there were many dimensions with structure that was clearly not noise. The projections in **Figure 7b** thus yield only a partial view. Subsequent quantification therefore employed all twelve dimensions.

**Figure 7.**
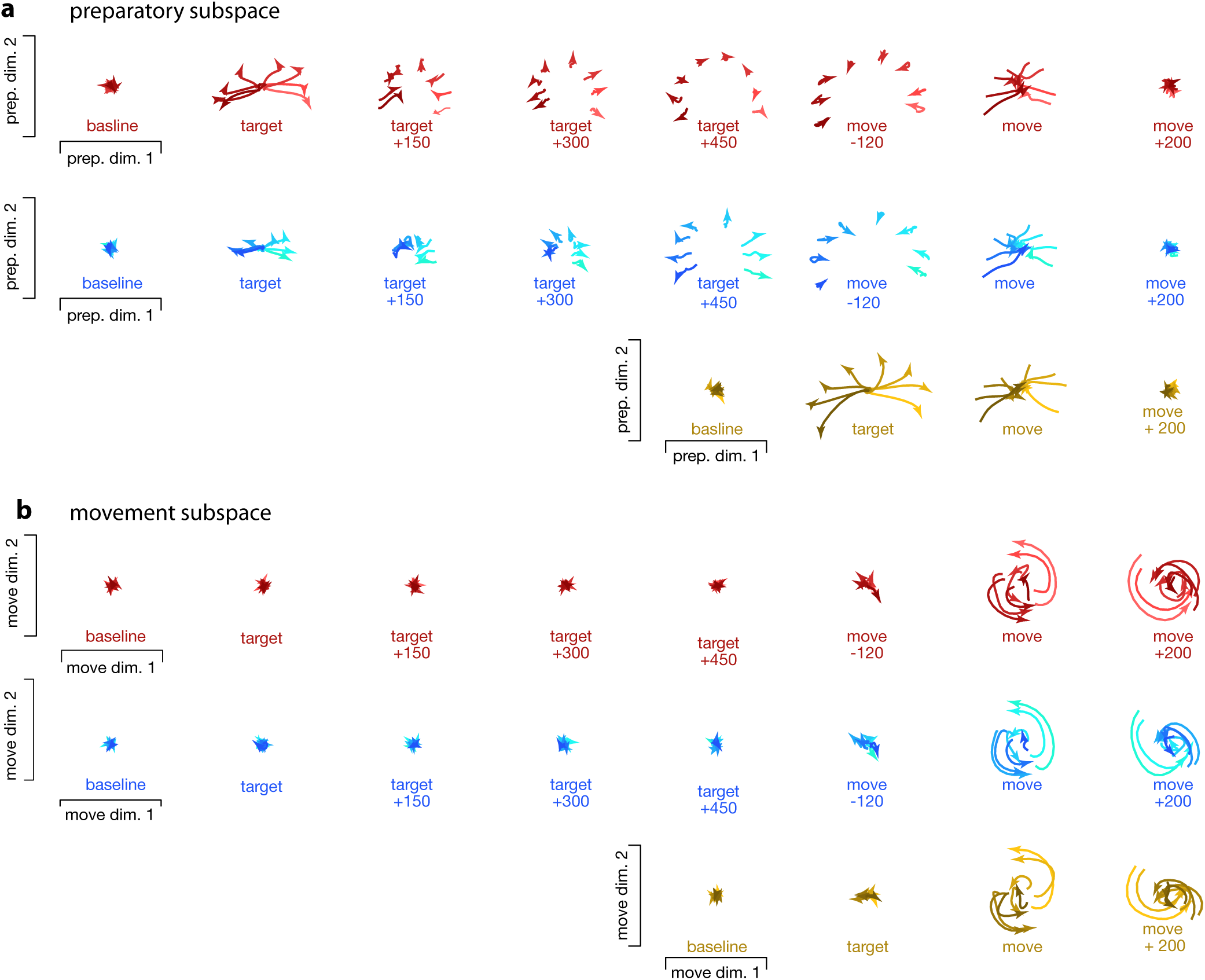
Snapshots of neural population state in the preparatory and movement subspaces for monkey Ba. Responses for cue-initiated *(red)*, self-initiated *(blue)* and quasi-automatic *(yellow)* contexts projected onto the top two preparatory dimensions (**a**) and top two movement dimensions (**b**). Those dimensions were found using the cue-initiated context data only. Trace colors correspond to target direction (same color scheme as in **Figure 3**). Each snapshot shows the neural state in that subspace, for all eight directions, across a 150 ms window. Snapshot labeled ‘baseline’ begins 150 ms before target onset. Snapshot labeled ‘target’ plots data starting at target onset. For the cue-initiated and self-initiated contexts, the subsequent three snapshots show activity in 150 ms increments, still aligned to target onset. Snapshots labeled ‘move −120’ start 120 ms before movement onset, with data aligned to movement onset. Subsequent panels begin at the indicated time.

There was a remarkable consistency, across contexts, in the patterns of the neural trajectories. The most notable differences across contexts regarded not the patterns *perse*, but the time-course of preparatory-subspace events. In the cue-initiated context, target onset prompted preparatory-subspace activity to become strongly selective for reach direction (*red traces* in **Fig 7a** separate upon target onset). The resulting pattern was sustained throughout the delay period, then collapsed near the time of movement onset. In the self-initiated context, target onset prompted a weaker separation of preparatory-subspace neural states (*blue traces* in **Fig 7a** separate less than *red traces* upon target onset). As time neared the onset of the self-initiated reach, the preparatory-subspace pattern became more robust until it was approximately as strong as that in the cue-initiated context. For the quasi-automatic context, the preparatory-subspace pattern was very shortlived: it grew rapidly following target onset then immediately collapsed prior to movement onset. However, while present, the preparatory-subspace pattern during the quasi-automatic context closely resembled that in the other two contexts (compare across contexts in the third-to-last column of **Fig 7a**). For example, the dependence of the neural state on reach direction was similar across contexts (*lighter / darker* traces indicate rightwards / leftwards movements).

Movement-subspace patterns were very similar across contexts, in both their pattern and their timing. Target onset produced essentially no separation of movement-subspace states for the cue-initiated or self-initiated contexts. This is consistent with finding that target onset produced little or no change in EMG activity. Movement-subspace states started to differentiate between reach directions ~110 ms before movement onset. This occurred at a similar time and in a similar way across contexts. During movement, the neural state evolved according to rotational dynamics, as previously reported^19^ and in a manner predicted by neural network models^32^. As for such models, rotational dynamics were present in a subset of dimensions; the dimensions shown here were chosen specifically to capture such dynamics for the cue-initiated context, and naturally captures similar dynamics for the other two contexts.

Comparing between subspaces reinforces and extends the results described in **Figure 6**. Across all contexts, preparatory-subspace activity always emerged before movement-subspace activity began. Preparatory-subspace activity and movement-subspace activity then showed considerable overlap: the former declined as the latter emerged. Just before and during that period of overlap, the pattern of preparatory subspace activity was similar across all three contexts. We explore this finding quantitatively below.

### Quantification of similarity across contexts

A central question of this study is whether similar movement-subspace events are preceded by similar preparatory subspace events. To quantify movement-subspace similarity, we measured the correlation (per dimension) between time-evolving patterns measured during a 150 ms window starting at movement onset. These correlations were high for all comparisons. Comparing the self-initiated and cue-initiated contexts, for the two dimensions shown in **Figure 7**, the correlation was 0.96 and 0.97 for monkey Ba and 0.97 and 0.95 for monkey Ax. Across all twelve movement dimensions, correlations ranged from 0.96 to 0.99 for monkey Ba and from 0.94 to 0.99 for monkey Ax. Correlations were similarly high when comparing the quasi-automatic and cue-initiated contexts. For the two dimensions shown in **Figure 7**, the correlation was 0.95 and 0.93 for monkey Ba and 0.91 and 0.97 for monkey Ax. Across all movement dimensions, correlations ranged from 0.88 to 0.99 for monkey Ba and from 0.90 to 0.98 for monkey Ax. These results agree with the similarity of movement-epoch activity across contexts observed at the individual-neuron level. How similar are the preceding patterns of preparatory subspace activity?

To address this question, we focused on the preparatory subspace state at a specific moment: 70 ms before movement onset. At this moment, movement-related activity is just starting to emerge. We have previously hypothesized that the preparatory state seeds movement-related dynamics^10,18,19,21,27,32^. Under this hypothesis, the preparatory state when movement-related activity begins is critical. This hypothesis thus predicts that the preparatory subspace state at that time should be very similar across contexts, given that subsequent patterns of movement-related activity are similar. We refer to that potentially critical preparatory state as the ‘final’ preparatory state; after that moment, movement-subspace activity becomes strong and preparatory-subspace activity declines to near-baseline levels.

The final preparatory state was similar across contexts. For each reach direction, the neural states across contexts formed a cluster (these are grouped via covariance ellipses in **Fig. 8a,b**). Clusters were quite tight for monkey Ba and somewhat less so for monkey Ax. To quantify the tightness of clustering – *i.e.*, the similarity of states across contexts – we computed the correlation, between contexts, of the set of preparatory-subspace states (one state per reach direction, with two dimensions describing each state). The correlation between cue-initiated and self-initiated contexts was 0.99 (95% c.i.: 0.98-0.99, p < 0.001) and 0.95 (95% c.i.: 0.86-0.98, p < 0.001) (monkey Ba and Ax). The correlation between cue-initiated and quasi-automatic contexts was 0.99 (95% c.i.: 0.97-1.0, p < 0.001) and 0.91 (95% c.i.: 0.76-0.97, p < 0.001). The correlation between self-initiated and quasi-automatic contexts was 0.99 (95% c.i.: 0.97-1.0, p < 0.001) and 0.92 (95% c.i.: 0.79-0.97, p< 0.001).

**Figure 8.**
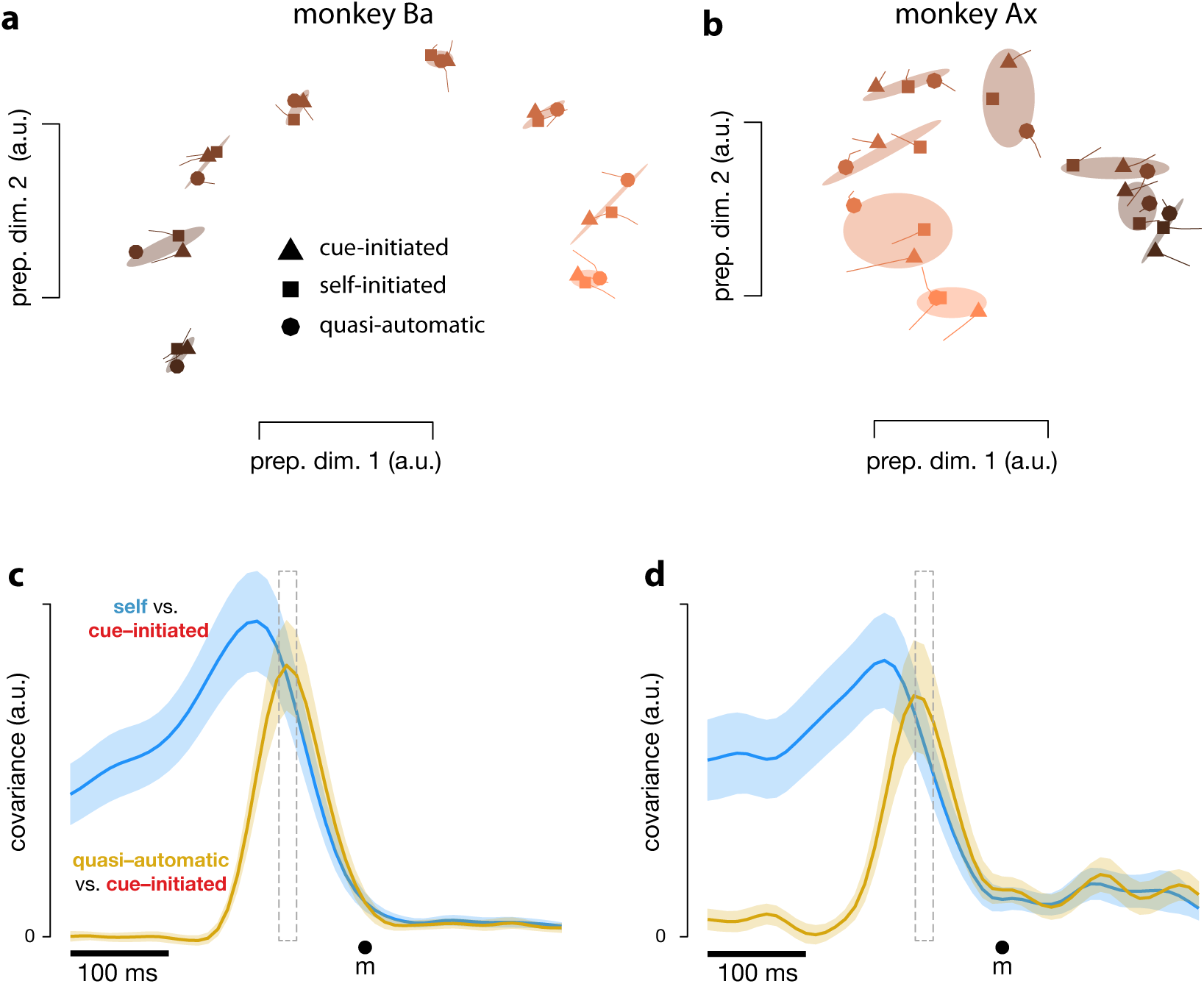
Preparatory subspace activity just before movement onset. **a)** Data for monkey Ba. Each marker denotes the neural state in a two-dimensional preparatory subspace: the two dimensions that captured the most variance, as in Fig. 7a. Markers indicate the state 70 ms before movement onset. Tails plot 20 ms of activity leading up to that time. The three shapes show states for the three contexts. Shaded regions plot a covariance ellipse for each triplet of states. A different symbol shade is used for each target direction *(light* for right, *dark* for left). **b)** As in **a** but for monkey Ax. **c**) Quantification of the time-course of the similarity in the pattern of preparatory states between contexts. Data is for monkey Ba. *Blue trace* plots the covariance between the preparatory pattern in the self-initiated context and that in the cue-initiated context. *Yellow trace* plots the covariance between the preparatory pattern in the quasi-automatic context and that in the cue-initiated context. The covariance is high when patterns are both strong and similar. Note that the units of the vertical scale are arbitrary (for reference, the correlation peaks close to one). Gray dashed window of time indicates the 20 ms time range (from 90 to 70 ms before movement onset) shown in **a** and **b**. The shaded regions denote the standard deviation of the sampling error (equivalent to the standard error) computed via bootstrap *(methods).*

To assess the time-course of similarity, at each time we computed the covariance, for a pair of contexts, between the neural states in the preparatory space. Covariance reflects both similarity and strength, and is thus expected to peak at a time when preparatory patterns are both similar and robust. When comparing the self-initiated and cue-initiated contexts (**Fig. 8c,d**; *blue*) covariance rose as movement approached, peaking 120 ms and 130 ms (monkey Ba and Ax) before movement onset. This is consistent with what can be observed in earlier figures: in the preparatory subspace, the pattern of states in the self-initiated context generally resembles that in the cue-initiated context, but is weaker until the time of movement onset nears.

When comparing the quasi-automatic and cue-initiated contexts (**Fig. 8c,d**; *blue*) covariance rose rapidly, peaking 80 ms and 90 ms (monkey Ba and Ax) before movement onset. These peaks occur just after activity in the movement-subspace first begins to change, which occurred 90 ms (monkey Ba) and 94 ms (monkey Ax) prior to movement onset in the quasi-automatic context. The narrowness of the peak underscores that the similarity in preparatory subspaces states was short-lived; it was high for only a few tens of milliseconds, around the time that movement-subspace activity was beginning to develop. Thus, while the pattern of preparatory subspace activity in the quasi-automatic context comes to closely match that in the cue-initiated context, this similarity occurs late (just as movement-subspace activity is developing) and is not sustained. This is consistent with a preparatory process that is observed across all contexts, but that unfolds very rapidly in the quasi-automatic context.

### Relative timing of movement-related events

Our subspace-based analysis method isolates a movement subspace that is, by construction, occupied during movement for the cue-initiated context. However, our method imposes no additional constraints on the timing of movement-subspace events: they could begin well before movement onset, at the time of movement onset, or after movement onset. We were particularly interested in the relationship between movement-subspace occupancy and the onset of muscle activity. Does movement-subspace occupancy occur with timing appropriate given a role in producing descending commands that cause muscle activity? For both movement-subspace occupancy and EMG, we assessed latency by measuring the moment when activity surpassed 10% of its peak. To minimize the impact of filtering on latency, these analyses employed a 10 ms Gaussian filter (rather than 20 ms for all other analyses) for both neural and EMG data.

Across monkeys and contexts, the movement subspace always became occupied just before the onset of changes in EMG, with an average latency of 21 ms. For comparison, the conduction delay from cortex to muscles, assessed via spike-triggered averages, can be as little as 6 ms from the time of a spike to the peak of the EMG response^33^. This delay would be slightly reduced (to ~4 ms for the lowest-latency neurons) when considering the beginning rather than the peak EMG response. Thus, activity in the movement subspace rises early enough to potentially account for the onset of muscle activity. This was consistently true across contexts, although with slight variability. The latency between the onset of movement-subspace activity and muscle activity was, for monkey Ba and Ax, 27 and 20 ms (cue-initiated), 33 and 22 ms (self-initiated) and 19 and 6 ms (quasi-automatic). These exact latencies should be interpreted with some caution: latencies are notoriously difficult to assess because high thresholds overestimate latency while low thresholds are sensitive to noise. Still, our best estimates indicate that, if cortico-motoneurons draw from movement-subspace activity, the onset of such activity occurs early enough to plausibly account for the onset of muscle activity.

## Discussion

### Is delay-period activity a reflection of motor preparation?

Early studies generally viewed delay-period activity as preparatory, but noted that directional selectivity often reverses between delay and movement epochs, suggesting a suppressive role^34^. Subsequent experiments revealed that delay-period and movement-related activity patterns typically differ^6,18,27,35^, ruling out the hypothesis that preparation involves a subthreshold version of movement-related activity. A different preparatory role for delay-period activity was suggested: serving as the initial state of a neural dynamical system whose evolution produces movement^10,18,21^. In support, one can directly observe that the phase and amplitude of movement-related dynamics flow from the state achieved during the delay^19^. Under this hypothesis, preparatory activity is a necessary precursor to movement-related activity. Yet a recent study yielded mixed evidence regarding the presence of a consistent preparatory state with and without a delay^22^. That mixed evidence highlighted the longstanding uncertainty regarding whether delay-period activity represents a true preparatory process, a facilitatory but non-obligatory process, or a suppressive process specific to an artificial imposed delay^23^.

Our results reveal that the neural process present during a delay-period is not specific to that situation, but is consistently observed in other contexts. This putatively preparatory process has the following properties. First, activity occupies a neural subspace orthogonal to that occupied during movement. Second, such activity consistently occurs before activity in the movement-related subspace. Third, regardless of the presence of an imposed delay period, the neural state in the preparatory subspace achieves a similar movement-specific state before movement onset. That similarity is maximal at the critical moment when movement-subspace activity is just beginning. These results essentially rule out the hypothesis that delay-period activity is primarily suppressive. The suppressive hypothesis cannot explain the presence of preparatory-subspace activity in the quasi-automatic context, or the rising profile of preparatory-subspace occupancy in the self-initiated context.

That said, our results do not prove that preparatory-subspace activity is preparatory – they only show that it follows the major predictions of that hypothesis. Proving that hypothesis would require specifically perturbing activity in that subspace and observing the impact on behavior – something not currently feasible. That said, it is known that a non-specific disruption of premotor cortex activity, at the end of the delay period, impacts RT in a manner consistent with disruption of a preparatory process^17^. Given that evidence and the present observations, we tentatively interpret preparatory subspace activity as preparatory and ask what conclusions might follow.

### Does preparation necessarily consume time?

Early behavioral investigations leveraged and supported the assumption that it takes considerable time to ‘plan’ or ‘specify’ the desired movement^2,5^. Influenced by this framework, subsequent physiology and modeling studies proposed that preparation involves the time-evolving strengthening and shaping of neural activity directly specifying movement parameters^9,36,37^. We have argued that movement is specified more implicitly, by achieving a preparatory state that seeds movement dynamics^18,19,21,27,32^, but found evidence that it takes time (100-200 ms) to consistently prepare^10^. Thus, a time-consuming preparatory process has often been considered to be a major determinant of RT. This traditional framework has enjoyed explanatory power, and has motivated successful comparisons of trial-to-trial RT variability with trial-to-trial variability of putatively preparatory activity^4,8,10,16^.

Yet there have been compelling recent arguments against the necessity of a time-consuming preparatory process^23,26^. The present study supports those arguments. RTs in the quasi-automatic context were on average 221 and 208 ms, and were frequently 170-200 ms on individual trials. These short RTs occur despite the inability to prepare in advance, and cannot be explained by anticipation: monkeys had no fore-knowledge of target direction and did not attempt to ‘jump the gun’. Given a delay of at least 50 ms for visual information to reach motor cortex, and an afferent delay of at least 75 ms (including the sizeable lag between muscle activity and movement onset), there cannot exist an obligatory preparatory process that necessarily takes 100-200 ms to complete. That conclusion is further supported by the neural data. In the quasi-automatic context, preparatory subspace activity lead movement-subspace activity by only ~40 ms, and the preparatory-subspace state came to match that in the cue-initiated context in ~70 ms. These findings rule out the idea of a slow, cognitive planning process that must complete before movement. These findings support our prior proposal that preparatory activity is necessary to seed movement-generating dynamics. However, the development of such activity can occur much faster than previously supposed.

Nevertheless, the influential idea that motor preparation tends to consumes time may have some merits. It may be that preparation often, or even typically, spans time. In the self-initiated context, putatively preparatory activity begins hundreds of milliseconds before movement-related activity. This raises a central question: if preparation can be fast, why is it ever extended? Why do monkeys not simply wait to prepare until just before movement onset? We can only speculate, but the ability to rapidly and consistently achieve the correct preparatory state may not be something that can be counted on in all real-world situations, especially for less familiar or more challenging movements. The motor system may thus have developed the conservative strategy of preparing in advance when possible, allowing time for errors to be corrected before movement generation begins^10,17^. We did indeed find that accuracy was slightly reduced in the quasi-automatic context, as would be expected if movement is sometimes triggered before preparation has fully converged on the appropriate preparatory state.

### Putatively preparatory and movement-related processes overlap

Preparatory subspace activity overlapped with movement subspace activity by slightly more than 100 ms. This overlap is consistent with (and indeed required by) the hypothesis that preparatory-subspace activity seeds movement-subspace dynamics. Aspects of the overlap explain a seeming discrepancy between our results and the recent finding of Ames et al.^22^ that the neural state for no-delay trials does not pass through the state achieved during the delay of long-delay trials. This might seem at odds with our finding that a consistent preparatory-subspace state is achieved across all contexts. In fact, our results are fully compatible. Preparatory-subspace activity in the quasi-automatic context achieves its maximal match with that in the cue-initiated context slightly after movement-subspace activity emerged. Thus, at the moment the match is achieved, the full neural state contains both preparation-related and movement-related contributions. The neural state at this moment will therefore not match that during the delay-period of the cue-initiated context, when there is no contribution from movement-related dimensions.

With this conflict resolved, the present results support and extend two key conclusions of Ames et al. First, the initial response to target onset can be similar with and without an imposed delay (compare the initial development of preparatory subspace activity between cue-initiated and quasi-automatic contexts in **Fig. 7**). However, this early response is unlikely to be an inevitable visual response: it is considerable weaker in the self-initiated context. Second, when under time pressure, the neural state does not momentarily pause at a stable state prior to the onset of movement-related activity (also see^8^). Indeed, in the quasi-automatic context, events are so compressed that preparatory-subspace activity is still developing as movement-subspace activity is beginning.

### Preparing versus deciding

Although our data argue against the conception of preparation as an intrinsically slow, cognitive process, they are quite consistent with the idea that slow cognitive processes influence preparatory activity. It is well established that preparatory activity in a variety of brain regions can reflect decisions regarding when or where to move^31,38-42^. Such decisions can sometimes unfold slowly or vacillate with time. In motor cortex, preparatory subspace activity may therefore sometimes evolve slowly simply because the overall movement goal is being decided slowly.

In the present study, the rising strength of pre-movement activity in the self-initiated context is somewhat reminiscent of the rise of choice-related activity in decision tasks. However, in the present case the target was always fully specified; target choice did not become more certain with time. Thus, strengthening pre-movement activity is unlikely to be related to target choice *perse*, and is more likely to reflect preparation to execute a choice that was clear from the outset (also see^31,43^). This suggests that having a clear movement goal does not necessarily mean that low-level preparatory processes are fully engaged. Whether or not preparatory activity develops may depend on whether it is reasonably likely that movement will be initiated soon. Consistent with this interpretation, studies that use a fixed, predictable delay typically find that delay-period activity ramps up with time (e.g.,^9^) while studies that use an unpredictable delay tend to find delay-period activity that reaches a rough plateau after a burst following target onset (e.g.,^10^).

Thus, the processes of deciding what to do, preparing to do it, and actually initiating, may occur with variable timing relative to one another. This hypothesis is potentially relevant to the finding that there exist neural events that are predictive of movement initiation, yet precede movement by more than the typical reaction time, and also precede self-report of the decision to initiate movement^43,44^. This is consistent with our finding that preparatory subspace activity, in the self-initiated context, develops hundreds of milliseconds before movement onset, and potentially before a definitive choice to execute movement has been made.

### Cortical involvement despite fast RTs

Reaching can involve very rapid, nearly involuntary corrections that are likely to have a subcortical contribution^25^. It has thus been suggested that entire movements may sometimes be produced subcortically, perhaps with minimal cortical involvement^26^. In particular, a loud, startling sound can release a pre-planned movement (the ‘StartReact’ phenomenon) with EMG-based RTs of 70-100 ms. While this short latency is due in part to the use of a highly salient auditory stimulus (which suffers less sensory delay than a visual stimulus), it also depends on the subcortically generated startle reflex. Given that the triggering impetus likely arises subcortically, it has been suggested that movement generation itself may not depend on cortical involvement. Yet a recent study argued against a reduction of cortical involvement in StartReact^45^. The authors instead interpreted StartReact as a subcortical triggering of movement-generating dynamics that span cortical and subcortical circuits in the same way as conventionally triggered movements. Our findings indicate that cortical events are not necessarily slow, and are thus compatible with this view. EMG activity in the quasi-automatic context could begin changing as early as 90 ms following target onset, with a mean of 150 ms and 130 ms for the two monkeys. This is not as fast as during StartReact, but is only slightly slower when one considers the additional sensory delay incurred by a visual stimulus (also note that in StartReact, movements are planned ahead of time while in the quasi-automatic context they are not). The fastest RTs in the quasi-automatic context are thus likely to be near the physiological limit. Yet we saw no evidence of reduced cortical involvement. Indeed, the patterns of cortical movement-related activity in the quasi-automatic context were very similar to those in the other two contexts. As in^45^, we do not suggest an absence of sub-cortical involvement, merely a conservation of cortical involvement.

The conservation of neural events across contexts in motor cortex should not be taken to imply that events other cortical or subcortical areas will be similarly conserved. Monkeys were clearly aware of the differences between contexts, and behaved appropriately. It must therefore be the case that some brain areas perform different computations in different contexts as necessary to initiate movement at the appropriate time. We have indeed observed that neural activity in the supplementary motor area differs across contexts not only during movement, but even before target onset (unpublished observations) and one suspects that this will be true of a variety of cortical and subcortical areas. However, motor cortex appears to be playing a more mechanical role: similar movements are driven by similar patterns of movement-related activity, following similar patterns of preparatory activity, across a broad range of timing constraints.

## Methods

### Subjects and task

Subjects were two adult male macaque monkeys *(Macaca mulatta)* aged 10 and 14 years and weighing 11 – 13 Kg at the time of the experiments. Daily fluid intake was regulated to maintain motivation to perform the task. All procedures were in accord with the US National Institutes of Health guidelines and were approved by the Columbia University Institutional Animal Care and Use Committee.

Subjects sat in a primate chair facing an LCD display and performed reaches with their right arm while their left arm was comfortably restrained. The timing of stimulus presentation was controlled using a photodetector (Thorlabs) to track individual frames on the display, such that the exact moment (within ~1 ms) of all events was known. This allows accurate assessment of reaction times and neural response latencies relative to visual events. Hand position was monitored using an infrared optical system (Polaris; Northern Digital) to track (~0.3 mm precision) a reflective bead temporarily affixed to the third and fourth digits. Each trial began when the monkey touched and held a central touch-point. Touch-point color indicated context (**Fig. 1**). After the touch-point was held for 450 – 550 ms (randomized) a colored 10 mm diameter disk (the target) appeared in one of eight possible locations radially arranged around the touch point. Target distance was 130 mm for cue and self-initiated contexts and 40 mm for the quasi-automatic context (**Fig. 1**). Trials for different contexts / directions were interleaved using a block-randomized design.

In the cue-initiated context, after a variable delay period (0-1000ms) the target suddenly grew in size to 30 mm and the central touch point simultaneously disappeared. These events served as the go-cue, instructing the monkey to make the movement. Reaches were successful if they were initiated within 500 ms of the go cue, had a duration < 500 ms, and landed within an 18 mm radius window centered on the target. Juice was delivered if the monkey held the target, with minimal hand motion, for 200 ms (this criterion was also shared across all three contexts).

In the self-initiated context, the target slowly and steadily grew in size, starting upon its appearance and ending when the reach began. Growth continued to a maximum size of 30 mm, which was achieved 1200 ms after target appearance (most reaches occurred before this time). The reward for a correct reach grew exponentially starting at 1 drop and achieved a maximum of 8 drops after 1200 ms. Monkeys were free to move as soon as the target appeared. However, monkeys essentially always waited longer in order to obtain larger rewards. In rare instances where no movement was detected 1500 ms after target onset, the trial was aborted and flagged as an error. Requirements for reach duration and accuracy were as for the cue-initiated context.

In the quasi-automatic context, the target moved radially away from the central touch-point at 25 cm / s. Motion began immediately upon target appearance. Target motion ended if a reach succeeded in bringing the hand to the target mid-flight. If the target was not intercepted (*e.g.*, if reach initiation was too slow) then the target continued moving until off the screen. Target speed and initial location (40 mm from the touch-point) were titrated, during training, such that the target was typically intercepted ~130 mm from the touch-point (the same location as the targets for the other two contexts). For successful interception, reaches had to land within an elliptical acceptance window (16 mm by 20 mm radius, with the long axis aligned with target motion). If the target was successfully intercepted, it grew in size to 30 mm and reward was delivered after the hold period.

Movement onset (and thus the RT) was measured based on hand speed: the magnitude of velocity in the plane of the movements (not including depth). We considered the time of movement onset to be the first moment when speed exceeded a threshold, set to 1% of average peak speed for that target location. To ensure this measure was robust on a given trial, we also measured ‘backwards’ from the moment of peak speed, and found the last moment when peak speed was below a threshold. On rare occasions, these ‘forwards’ and ‘backwards’ measurements disagreed (*e.g.*, due to an unusual velocity profile) in which case the time of movement onset was considered undefined and the RT was not analyzed.

### Neural and muscle recordings

After subjects became proficient in the task, we performed sterile surgery to implant a head restraint. At the same time, we implanted a recording chamber centered over the arm area of primary motor cortex (M1) and the dorsal premotor cortex (PMd) of the left hemisphere. Chamber positioning was guided by structural magnetic resonance images taken shortly before implantation. We used intracortical microstimulation to confirm that our recordings were from the forelimb region of motor cortex (biphasic pulses, cathodal leading, 250μS pulse width delivered at 333 Hz for a total duration of 50 ms). Microstimulation typically evoked contractions of the shoulder and upper-arm muscles, at currents from 5 ?A – 60 ?A depending on the location and cortical layer. We recorded single-neuron responses using traditional tungsten electrodes (FHC) or one or more silicon linear-array electrodes (V-probes; Plexon) lowered into cortex using a motorized microdrive. For tungsten-electrode recordings, spikes were sorted online using a window discriminator (Blackrock Microsystems). For linear-array recordings spikes were sorted offline (Plexon Offline Sorter). We recorded all well isolated task-responsive neurons; no attempt was made to screen for neuronal selectivity for reach direction or any other response property. Spikes were smoothed with a Gaussian kernel with standard deviation of 20 ms and averaged across trials to produce peri-stimulus time histograms. For measurements of latency, we used a 10 ms Gaussian to minimize the impact of filtering on latency.

We recorded electromyogram (EMG) activity using intramuscular electrodes from the following muscles: lower and upper aspects of the trapezius, medial, lateral and anterior aspects of the deltoid, medial and outer aspects of the biceps, brachialis, pectoralis and latismus dorsi. The triceps were minimally active and were not recorded. EMG signals were bandpass filtered (10 – 500 Hz), digitized at 1kHz, rectified, smoothed with a Gaussian kernel with standard deviation of 20 ms, and averaged across trials to produce peri-stimulus time histograms.

### Data pre-processing prior to population analyses

As in our previous work, we employed two pre-processing steps ^19^. First, the responses of each neuron were soft-normalized so that neurons with high firing rates had approximately unity firing-rate range (normalization factor = firing rate range+5). This step ensures that subsequent dimensionality reduction (see below) captures the response of all neurons, rather than a handful of high firing-rate neurons. Second, the responses for each neuron were mean-centered at each time as follows: we calculated the mean activity across all conditions of each neuron at each time point, and subtracted this mean activity from each condition’s response. This step ensures that dimensionality reduction focuses on dimensions where responses are selective across conditions, rather than dimensions where activity varies in a similar fashion across all conditions^46^.

### Identifying preparatory and movement dimensions

We recently developed a method that leverages the finding that neural responses in the delay-epoch are nearly orthogonal to responses in the movement epoch^27^. The method identifies one set of preparatory dimensions and an orthogonal set of movement dimensions. Briefly, we define two matrices based on data from the cue-initiated context only: *P* ∈ ℝ^*N* × *CT*^ which holds preparatory epoch responses and *M* ∈ ℝ^*N* × *CT*^ which holds movement-epoch neural responses. *N* is the number of neurons recorded, *C* is the number of reach directions and *T* is the number of time points. The method seeks to find a set of preparatory dimensions, 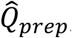, that maximally capture the percentage of variance of *P* and an orthogonal set of movement dimensions, 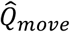, that maximally capture the percentage of variance of *M*. We compute the preparatory and movement-epoch covariance matrices *C*_*prep*_ = *cov*(*P*) and *C*_*move*_ = *cov*(*M*) and optimize the following objective function:

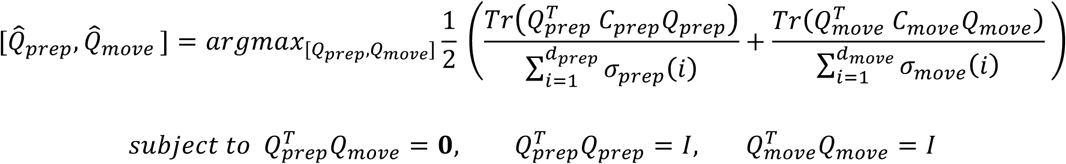

where *σ*_*prep*_(*i*)is the *i*^*th*^ singular value of *C*_*move*_. *Q*_*prep*_ and *Q*_*move*_ are the bases for the preparatory and movement subspaces respectively and *Tr*(·) is the matrix trace operator. The term 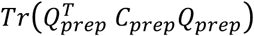 reflects the preparatory-epoch data variance captured by the preparatory subspace, and 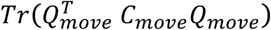 reflects the movement-epoch data variance captured by the movement subspace. We chose the dimensionality of *Q*_*prep*_ to be 12 (i.e., *Q*_*prep*_ ∈ ℝ^*N × 12*^), which captured ~ 80% of preparatory-epoch variance (the remaining variance had very little structure and appeared to be primarily sampling noise). Similarly, we chose the dimensionality of *Q*_*move*_ to be 12, which captured ~ 85% of movement-epoch variance. Results were robust with respect to the choice of dimensionality. The optimization objective is normalized (by the singular values) to be insensitive to the relative dimensionality and amount of response variance in the two subspaces. This normalization is particularly important in our case since movement activity is stronger and typically has higher dimensionality than the preparatory activity. For visualization, we need to choose two dimensions spanned by *Q*_*prep*_and *Q*_*move*_to define the plotted projections (*e.g.*, in **Fig. 7**). For *Q*_*prep*_we chose the basis so that the top two dimensions captured the most variance (with all others ranked accordingly). For *Q*_*move*_ the basis was chosen using the jPCA method^19^ to capture movement-related oscillatory activity patterns.

### Projections and reconstructions

For a given time *t* and for condition *θ*, the projection of the population response onto the *k^th^* preparatory dimension is simply a weighted sum of all single-neuron responses: 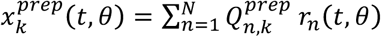 where 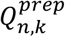 is the element in the *n*^*th*^ row and *k*^*th*^ column of *Q*_*prep*_ (see previous section) and *r*_*n*_(*t, θ*) is the response of the *n^th^* neuron. This is illustrated in Figure 4, where the orange weights *w* are taken from *Q*_*prep*_. The projection onto each movement dimension is defined analogously. The response of a given neuron can then be reconstructed as: 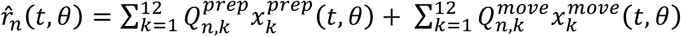. The two sums on the right-hand side are the preparatory and movement contributions respectively, which we can term 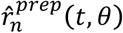 and 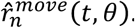. These are the preparatory and movement patterns for neuron *n*.

### Subspace occupancy

Our measurement of subspace occupancy is equivalent to the variance explained metric often computed in the context of principal component analysis. For the preparatory subspace, occupancy was computed as: 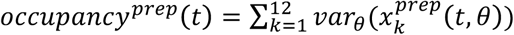 where ^*var*^*θ* indicates taking the variance across conditions (i.e., directions). Because dimensions are orthonormal this is equivalent to computing the variance for each neuron’s preparatory pattern and then summing: 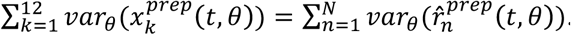. Movement subspace occupancy was defined analogously. To estimate the sampling error of the subspace occupancy we used a bootstrap procedure. We created 1000 surrogate populations by redrawing with replacement from the original population. We computed the subspace occupancy for each surrogate population, and for each time computed the standard deviation across these 1000 measures.

### Latency of physiological events

To measure the latency of preparatory-and movement-subspace occupancy, we filtered the spike trains of all neurons using a Gaussian kernel with 10 ms standard deviation. We recomputed the preparatory and movement dimensions using these data and calculated the subspace occupancy as before. We measured the latency as the first moment in time in which occupancy reached 10% of peak occupancy.

Similarly, to calculate the latency of the EMG with respect to movement onset, we filtered EMG activity of all muscles using a Gaussian kernel with a 10 ms standard deviation. We then performed PCA on the EMG activity for each context separately and projected the corresponding EMG responses onto the first PC. We measured the latency as the first moment in which activity in the first PC reached 10% of peak activity.

## Acknowledgments

We thank Yanina Pavlova and Sean Perkins for technical support. This work was supported by the Sloan Foundation, the Simons Foundation (SCGB#325171 and SCGB#325233), the Grossman Center for the Statistics of Mind, the McKnight Foundation, an NIH Directors New Innovator Award (DP2 NS083037), NIH/NSF CRCNS R01NS100066, the Kavli Foundation, a Klingenstein-Simons Fellowship, the Searle Scholars Program, an NIH Postdoctoral Fellowship (F32 NS092350) and the Gatsby Charitable Trust. We thank Andrew Zimnik for helpful discussions and performing some analyses of behavior.

### Author Contributions

A.H.L. and M.M.C. designed the experiments. A.H.L recorded and analyzed the data. G.F.E. and J.P.C. developed and implemented the method for segregating preparatory and movement subspaces. All authors contributed to analysis choices and interpretation. A.H.L. and M.M.C. wrote the manuscript with help from G.F.E. and J.P.C.

### Competing Financial Interests

The authors declare no competing financial interests.

---

**Supplementary Figure 1.**
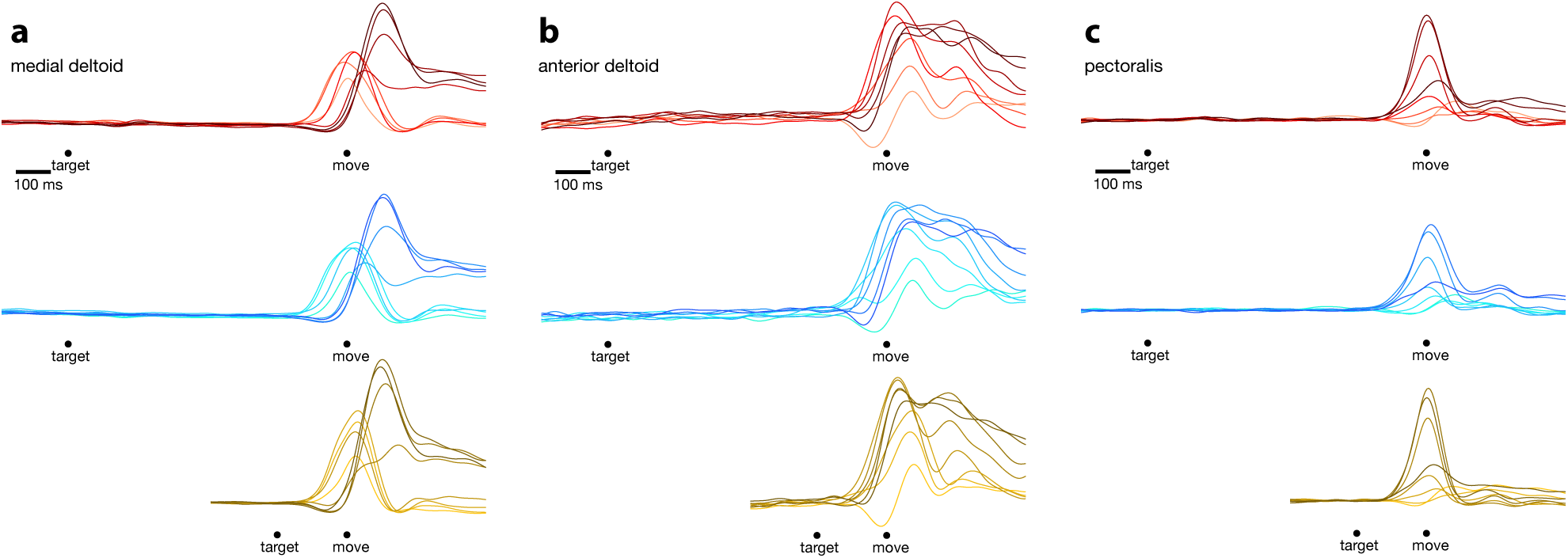
Responses from three muscles of the upper arm. Each column shows responses of a single muscle for the three initiation contexts. Each trace denotes the trial-averaged firing rate for one reach direction. Individual EMG records were filtered and rectified before smoothing with a Gaussian and averaging. Alignment of individual trials to task events is as described in Fig. 3 of the main text.

**Supplementary Figure 2.**
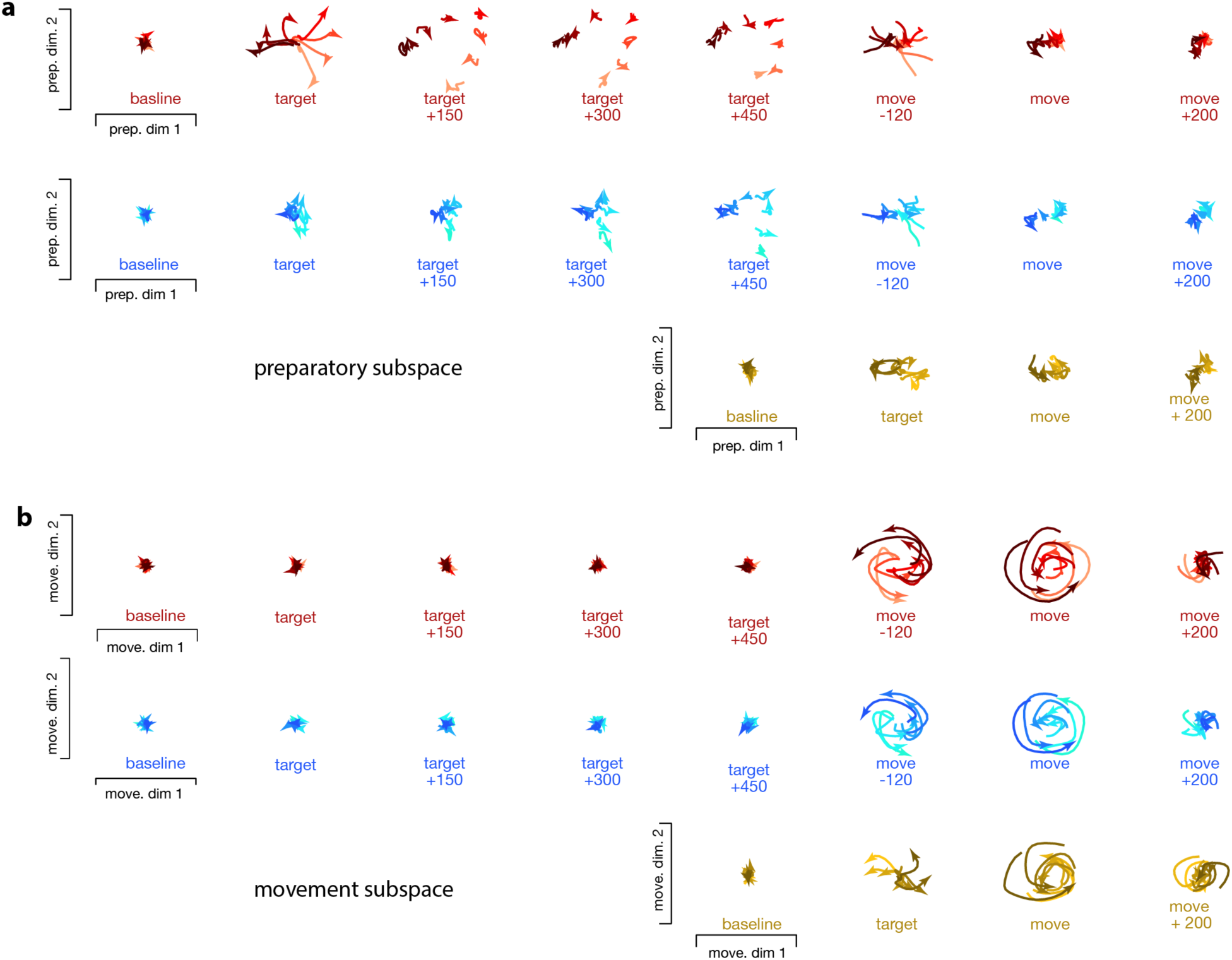
Snapshots of neural population state in the preparatory and movement subspaces for monkey Ax. All plotting conventions are as in Figure 7 of the main text.

